# Transcriptional and morpho-physiological responses of *Marchantia polymorpha* upon phosphate starvation

**DOI:** 10.1101/2020.09.16.300814

**Authors:** Felix Rico-Reséndiz, Sergio Alan Cervantes-Pérez, Annie Espinal-Centeno, Melissa Dipp-Álvarez, Araceli Oropeza-Aburto, Enrique Hurtado-Bautista, Andres Cruz-Hernández, John L. Bowman, Kimitsune Ishizaki, Mario A. Arteaga-Vázquez, Luis Herrera-Estrella, Alfredo Cruz-Ramírez

## Abstract

Phosphate (Pi) is a pivotal nutrient that constraints plant development and productivity in natural ecosystems. Land colonization by plants, more than 470 million years ago, evolved adaptive mechanisms to conquer Pi-scarce environments. However, little is known about the molecular basis underlying such adaptations at early branches of plant phylogeny. To shed light on how early divergent plants respond to Pi limitation, we analyzed the morpho-physiological and transcriptional dynamics of *Marchantia polymorpha* upon Pi starvation. Our phylogenomic analysis highlights some gene networks present since the Chlorophytes and others established in the Streptophytes (eg. PHR1-SPX1 and STOP1-ALMT1, respectively). At the morpho-physiological level, the response is characterized by the induction of phosphatase activity, media acidification, accumulation of auronidins, reduction of internal Pi concentration and developmental modifications of rhizoids. The transcriptional response involves the induction of Mp*PHR1*, Pi transporters, lipid turnover enzymes and Mp*MYB14*, an essential transcription factor for auronidins biosynthesis. Mp*STOP2* up-regulation correlates with expression changes in genes related to organic acid biosynthesis and transport, suggesting preference for citrate exudation. Analysis of MpPHR1 binding sequences (P1BS) shows enrichment of this *cis* regulatory element in differentially expressed genes. Our study unravels the strategies, at diverse levels of organization, exerted by *M. polymorpha* to cope with low Pi availability.

**Significance Statement:** This study unravels the transcriptional and morphophysiological mechanisms executed by the non-vascular, and rootless, plant *Marchantia polymorpha* upon phosphate starvation conditions. The findings in this study shed light on the mechanisms that early land plants may have developed for the conquest of substrates poor in available phosphate, some of which are still conserved by current-day plants. Moreover, our results open several working hypotheses and novel perspectives for the study of Pi-starvation responses along plant evolution.

## Introduction

The colonization of land by plants was a complex ecological, evolutionary and developmental (eco-evo-devo) process that occurred around 470 million years ago (Sanderson, 2003; Delwiche and Cooper, 2015). Several developmental innovations, which allowed early plants to adapt and survive on terrestrial environments, have been deeply reviewed elsewhere (Pires and Dolan, 2012; Ishizaki, 2017; Dipp-Alvarez and Cruz-Ramírez, 2019). However, little is known about the molecular innovations that allowed plants to thrive in conditions of low nutrient availability. The molecular mechanisms underlying the morphological and physiological strategies to forage low available nutrients, have been mostly described in diverse plant root systems. This is the case for most studies focused on describing the adaptation of plants to phosphate starvation (Lopez-Bucio et al., 2003., Puga et al., 2017; Gutiérrez-Alanís et al., 2018). Phosphorus is an essential macronutrient for proper plant development, which can only be acquired as ortho-phosphate (Pi) by plant roots (Raghothama, 1999). In natural ecosystems, Pi is generally distributed at low concentrations due to the rapid formation of inorganic complexes with Ca^+2^ or Al^+3^ and Fe^+3^, under alkaline or acid conditions, respectively (Bieleski, 1973; Marschner, 2011). This phenomenon causes that up to 70% of global agricultural areas display a limited Pi availability, affecting both plant development and crop productivity (Lynch, 1995). To enhance Pi uptake and recycling, diverse molecular mechanisms underlying plant morphological and physiological strategies are induced (Puga et al., 2017; Gutiérrez-Alanís et al., 2018). In *Arabidopsis thaliana* these strategies have been characterized and classified in local and systemic responses (Thibaud et al., 2010). The local response largely depends on the contact of the primary root tip with a Pi-depleted substratum, whose final outcome is the full differentiation of the primary root meristem and the triggering of lateral root development (Svistoonoff et al., 2007). One of the early events that takes place in the local response of Arabidopsis to this stress, involves the accumulation of the protein SENSITIVE TO PROTON RHIZOTOXICITY1 (AtSTOP1), a zinc finger transcription factor (TF) (Balzergue et al., 2017). Once in the nucleus, AtSTOP1 promotes the expression of several genes including the *ALUMINIUM ACTIVATED MALATE TRANSPORTER 1* (*AtALMT1*), which encodes a protein that enhances malic acid efflux to the apoplast (Balzergue et al., 2017; Mora-Macias et al., 2017). The malate efflux contributes significantly to the modification of the iron distribution and impacts directly the developmental program that defines the Arabidopsis root system architecture (RSA) under low Pi availability, Al^+3^ toxicity and acid pH (Balzergue et al., 2017; Mora-Macias et al., 2017). Also, AtSTOP1 induces the expression of *REGULATION OF ALMT1 EXPRESSION 1* (*AtRAE1*), which encodes an F-Box protein that ubiquitinates STOP1, inducing its degradation via the 26S proteasome pathway (Zhang et al., 2019). Additionally, exudation of organic acids (OAs) releases Pi from insoluble forms with Al^+3^ or Fe^+3^, improving the Pi assimilation in low pH conditions (Kochian et al., 2015). However, the post-translational regulatory mechanisms that control AtSTOP1 accumulation, in both high and low Pi conditions, have still not fully understood. Also, as part of the local response, a multicopper oxidase encoded by the *LOW PHOSPHATE ROOT 1* (*AtLPR1*) gene, changes the iron oxidation state from Fe^2+^ to Fe^+3^ in the root tip apoplast under low Pi conditions, this phenomenon correlates with callose accumulation and triggers a molecular cascade that leads to the full differentiation of the root apical meristem (RAM) (Müller et al., 2015). Furthermore, iron redistribution and its redox fluctuations, coupled to reactive oxygen species (ROS) signaling, promote the expression of *CLAVATA3/ENDOSPERM SURROUNDING REGION 14* (*AtCLE14*) gene which codes for a peptide that triggers the full differentiation of the RAM. The AtCLE14 peptide is sensed by *PEPTIDE RECEPTOR 2* (*AtPEPR2*) and *CLAVATA 2* (*AtCLV2*) receptors which, in turn, may act negatively on POLTERGEIST (AtPOL) and POLTERGEIST-LIKE 1 (AtPLL1), two protein phosphatases that are needed for the proper expression of *SCARECROW* (*AtSCR*) and *SHORT ROOT* (*AtSHR*), both essential factors for root stem cell niche (RSCN) maintenance (Song et al., 2008; Gutiérrez-Alanís et al., 2017).

The systemic response to low Pi availability is controlled by the internal concentration of inositol polyphosphate (IPP) and it is mainly modulated by PHOSPHATE STARVATION RESPONSE 1 (AtPHR1), a MYB-CC TF that activates the expression of several genes involved in the transport, scavenging and recycling of Pi (Rubio et al., 2001; Wild et al., 2016). This TF was first characterized in the Chlorophyte algae *Chlamydomonas reinhardtii*, under the name of PHOSPHORUS STARVATION RESPONSE 1 (CrPSR1) (Shimogawara et al., 1999; Wykoff et al., 1999). In *A. thaliana*, PHR1 controls the transcription of high affinity Pi transporters to improve nutrient assimilation under Pi deprived conditions. For example, AtPHR1 promotes the expression of *PHOSPHATE TRANSPORTER 1* (*AtPHT1*) to enhance Pi transport across the plasma membrane (PM) of epidermal and root cap cells (Nussaume et al., 2011; Kanno et al., 2016). AtPHR1 induces also the expression of *AtPHT2, AtPHT4* and *AtPHT3* genes, which encode transporters involved in intracellular Pi redistribution from cytosol to chloroplast and mitochondria (Daram et al., 1999; Versaw and Harrison 2002; Guo et al., 2008; Aguo et al., 2008B; Takabatake et al., 1999; Jia et al., 2015). The *VACUOLAR PHOSPHATE TRANSPORT 1* (*AtVPT1*) and *VACUOLAR PHOSPHATE EFFLUX TRANSPORTER 1* (*AtVPE1*) genes encoding transporters involved in Pi efflux from vacuole to cytosol and vice versa (Liu et al., 2015; Xu et al., 2019). Another aspect of the systemic response controlled by AtPHR1 is the recycling of internal Pi from phospholipids and nucleic acids (Essigmann et al., 1998; Cruz-Ramirez et al., 2006). Pi-deprived plants re-organize the PM composition to maintain cell integrity, replacing membrane phospholipids with non-phosphorus lipids such as sulfoquinovosyldiacylglycerol and digalactosyldiacylglycerol (Essigmann et al., 1998; Andersson et al., 2003; Cruz-Ramirez et al., 2006). Under low Pi conditions, AtPHR1 promotes the transcription of key genes involved in the lipid turnover pathway such as *AtSQD1* and *AtPLDz2* (Andersson et al., 2003; Cruz-Ramirez et al., 2006). Another key response controlled by AtPHR1 is the induction of *PURPLE ACID PHOSPHATASES* (*AtPAPs*) and *RIBONUCLEASES LIKE 1* (*AtRNS1*) genes, that encode enzymes that are secreted to the rhizosphere and participate in the release of Pi from the organic pool (Bariola et al., 1994; Li et al., 2002). The activity of AtPHR1 is negatively regulated by proteins containing the SIG1-PHO81-XPR1 (SPX) domain (encoded by *AtSPX1, AtSPX2, AtSPX3* and *AtSPX4* genes), such post-translational regulation of AtPHR1 occurs as a function of internal inositol polyphosphate (IPP) levels (Puga et al., 2014; Wang et al., 2014; Lv et al., 2014; Osorio et al., 2019). In high Pi conditions, AtSPX4 negatively regulates AtPHR1, through protein-protein interaction, at the cytoplasm (Lv et al., 2014; Osorio et al., 2019), while at low Pi conditions, AtPHR1 promotes the expression of *AtSPX1/2*, that act as repressors of PHR1 activity in the nucleus (Puga et al., 2014; Wang et al., 2014; Wild et al., 2016).

Most of the studies focused on characterizing the response of plants to Pi scarcity employed *A. thaliana’s* root as a model. The root is an essential organ for water and nutrient uptake in diverse clades of the Viridiplantae kingdom, however we should consider that the root is an innovation which occurred during land plant evolution millions of years later than the developmental and physiological mechanisms which allowed plants to thrive in soils with limited water and nutrients.

Pioneer studies on the response of *M. polymorpha* to Pi limitation described changes in rhizoid abundance, red thallus pigmentation and precocious formation of gemmae cups (Shull C, 1925; Voth & Hamner, 1940; Voth, 1941). Voth and Hamner (1940) suggested that some segments of thallus tips grow well under low Pi conditions, maybe due to a storage of nutrients which occurred previous to the transference to low Pi conditions. Such studies pointed already to the existence of a molecular program in liverworts to deal with Pi scarcity. However, the molecular and physiological mechanisms of Marchantia to adapt to, and develop in, an environment with low Pi levels remain obscure. In an attempt to shed light on the existence and conservation of such adaptive mechanisms in a rootless early divergent land plant, in this study we describe the morpho-physiological changes, and the underlying transcriptional profile, of *Marchantia polymorpha* in response to low Pi availability.

## Results

### 1) Evolutionary landscape of Pi starvation-induced genes across the plant kingdom

In the last decades, diverse studies revealed how key genetic networks orchestrate adaptive mechanisms of plants to cope with Pi-limited conditions (Puga et al., 2017; Gutiérrez-Alanís et al., 2018). However, little is known about how these strategies evolved along the early divergent lineages of the plant kingdom. To explore the conservation and divergence of such genetic mechanisms, we performed a survey in key representative genomes distributed across plant phylogeny (Figure 1a). Putative homologs were searched by sequence homology and phylogenetic reconstructions, which allowed us to identify the evolutionary history of genes related to Pi sensing, organic acid exudation, lipid metabolism, and Pi transport (*Figure 1a*).

**Figure 1.**
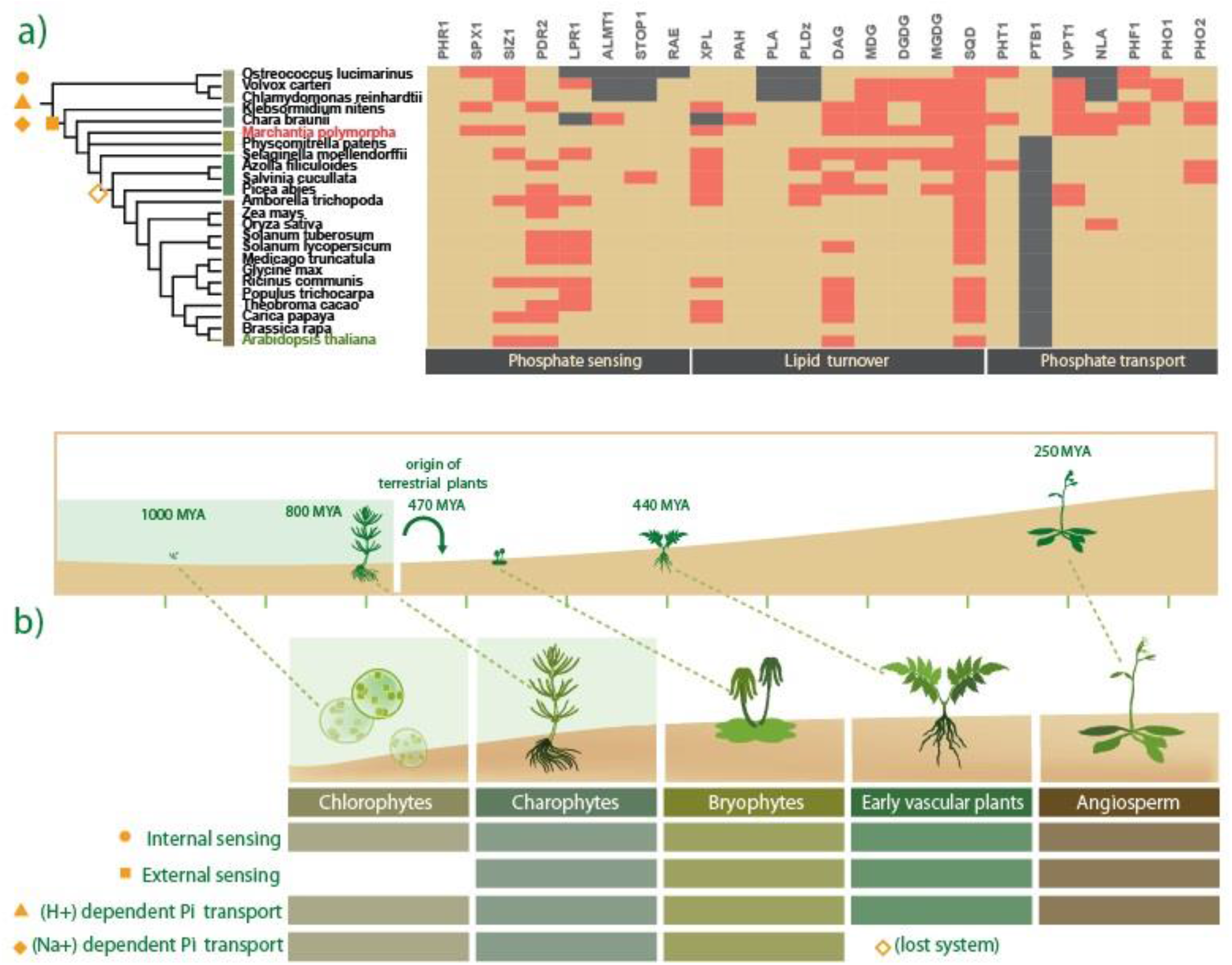
Evolution of phosphate starvation responses in land plants. a) Phylogenetic tree containing 24 representative genomes sampled across plant kingdom, the cladogram was taken from the NCBI taxonomy browser (https://www.ncbi.nlm.nih.gov/Taxonomy/CommonTree/www.cmt.cgi). The heatmap shows a comparative conservation analysis of key genes involved in Pi sensing, lipid turnover and Pi transport. Red-colored rectangles represent single copy genes, beige-colored rectangles mean that for such proteins there are 2 or more genes, and grey-colored mean that no gene coding for such proteins was identified through our approach (see methods for details). a) Showed a land plant evolution timeline. A close up highlight the morphological differences between chlorophytes, charophytes, bryophytes, gymnosperms and angiosperms. Color rectangles showed the conservation for key mechanisms to cope with Pi starvation, such as internal sensing network, external sensing network, (Na+) and (H+) dependent Pi transporters.

In the Pi sensing category, putative genes encoding the internal Pi sensing module *PHR1-SPX1* were found in all clades analyzed, from Chlorophyte algae to land plants (*Figure 1b*). In contrast, in the local Pi sensing genetic network composed of the *STOP1, ALMT1, LPR1, PDR2* and *RAE1* genes, some members were absent in the Chlorophyte lineage, while each was present in all the Streptophyte lineage (*Figure 1b*). Interestingly, the genome of *Ostreococcus lucimarinus* lacks 4 of the 5 genes related to local sensing, but the other Chlorophyte genomes analyzed (*Volvox carteri* and *Chlamydomonas reinhardtii*) possess putative homologs genes of *PDR2, LPR1*, and *RAE1* (*Figure 1a*). The absence of the STOP1-ALMT1 module in the Chlorophyte sampled genomes was evident, whereas in charophyte algae (*Chara Braunii* and *Klebsormidium nitens*) putative homologs of STOP1 and ALMT1 were present (*Figure 1a*). Our evolutionary analyses highlighted the STOP1-ALMT1 module as a streptophyte innovation, while the internal Pi sensing module PHR1-SPX1 was present since early green algae lineages (*Figure 1b*). Moreover, the putative homologs implicated in lipid turnover, such as XPL, DAG, MGD, DGDG, MGDG, and *SQD* were highly conserved in all the lineages investigated (*Figure 1a*). In addition, we explored Pi transport mechanisms and found putative homologs of *PHT1* and *PHF1* in all the species analyzed (*Figure 1a*), but the putative homologs of *NLA* were only present in the streptophyte lineage (*Figure 1a*). We found that *PTB* genes were present exclusively in both green algae lineages (Chlorophyte and Charophyte) and bryophyte (*Figure 1b*) as previously reported (Bonnot et al., 2017). Moreover, we found homologs of *VPT* in all sampled genomes, with the exception of *O. lucimarinus* (*Figure 1a*). The last transporter family analyzed was the AtPHO1, we found putative homologs in all species analyzed in this study (*Figure 1a*).

### 2) *M. polymorpha* exhibits low genetic redundancy in key Pi-responsive regulatory networks

*M. polymorpha* provides a useful model system due to its well characterized morphology, anatomy complexity (Shimamura et al., 2016), and its low genetic redundancy (Bowman et al., 2017). Our presence-absence analysis of putative homologs related to PSR (*Figure 1a*), led us to uncover a low genetic redundancy for key genetic networks in *M. polymorpha*, as compared to other land plants (*red rectangles in figure 1a*). For example, the internal Pi sensing module PHR1-SPX1 contains 11 and 4 members of each family in *A. thaliana*, respectively. While in the *M. polymorpha*, we identified only 3 genes encoding putative PHR/PSR-related MYB-CC TFs (*Figure 1a*). Our phylogenetic reconstruction placed MpPHR1 (Mapoly0003s0147) and MpPHR2 (Mapoly0098s0044) close to AtPHR6, in contrast MpPHR3 (Mapoly0115s0048) resides in another subclade with AtPHR8/9/12 (*Supplementary Figure S3*). *AtPHR1* and *AtPHL1* form an independent clade which includes protein sequences from the early vascular plant *S. moellendorffii* up to *A. thaliana*, without bryophyte or green algae sequences. While a single gene encoding a SPX-class 1 member was found in *M. polymorpha*, Mp*SPX* (Mapoly0002s0123) is phylogenetically closest to AtSPX4 (*Supplementary Figure S2*).

In the lipid turnover category, there are two *AtDAG* genes, two *AtPAH*, two *AtSQDs*, two *AtDGDGs*, three *AtMGDGs*, three *AtMGDs* and two *AtPLDzs* in the *A. thaliana* genome, while in *M. polymorpha* genome, our analysis retrieved single homologs for *PAH, MGD, DGDG, MGDG* and *SQD*, (*Figure 1a*). In the opposite sense, we found nine homologs for *PLDz* and seven for *PLA*. Our phylogenetic reconstructions showed that Mp*PAH* (Mapoly0089s0039), Mp*DAG* (Mapoly0022s0098), Mp*MGD* (Mapoly0023s0061), Mp*MGDG* (Mapoly0023s0061) and Mp*SQD* (Mapoly0010s0049) sequences were in a clade with putative orthologs from *P. patens* and *S. moellendorffii* (*Supplementary Figure S11, S14, S15, S17 and S18*). While the phylogenetic tree of DGDG family consists of two clades, one for the *AtDGDG1* and other for *AtDGDG2*, the latter includes the single Mp*DGDG* sequence (*Supplementary Figure S16*). Moreover, we found a single vacuolar Pi transporter in *M. polymorpha, MpVPT* (Mapoly0094s0020), while the *A. thaliana* genome has 3 loci (Liu et al., 2015). Phylogenetic reconstruction of *VPT-like* proteins showed that Mp*VPT* (Mapoly0094s0020) is in a clade with putative homologs from other bryophytes (*Supplementary Figure S20*). Finally, the *M. polymorpha* genome has a single homolog, Mp*NLA* (Mapoly0044s0127), while *A. thaliana* possesses two copies where the phylogenetic reconstruction places Mp*NLA* close to other bryophyte sequences (*Supplementary Figure S21*).

### 3) Phenotypical and physiological impacts of Pi availability on thallus development

In natural ecosystems, fluctuations of Pi concentrations in the soil drive fine-tuned mechanisms that allow plants to cope with limiting Pi conditions (Puga et al., 2017; Gutiérrez-Alanís et al., 2018). For example, root morphological modifications and physiological adaptations which respond locally to Pi availability, have been associated with enhanced Pi assimilation (Peret et al., 2011). However little is known about the phenotypic plasticity upon Pi starvation in plants without roots, especially in liverworts such as *M. polymorpha*. To characterize the morpho-physiological responses to low Pi availability during thalli development, gemmae were grown on agar media supplemented with different concentrations of Pi (0, 10, 25, 50, and 500 μM) and their phenotypes were recorded at multiple time points (7, 14 and 21 days) (*Supplementary figure S25*). Allowing us to determine Pi concentrations for high (+Pi) and low (-Pi) availability. The production of a red pigment, most probably auronidin accumulation (Albert et al., 2018; Berland et al., 2019), and changes in rhizoid development were observed at 0, 10 and 25 μM of Pi (*Supplementary figure S25*). Thus, taken in consideration that thallus development was severely affected at 0 μM and 25 μM was the highest Pi concentration that induced phenotypic changes, we selected 10 μM as a low Pi availability condition. We defined 500 μM of Pi as high availability condition, since at this concentration thalli neither showed accumulation of auronidin nor differences in rhizoid development (*Supplementary figure S25*). After registering the alteration on rhizoid development under −Pi conditions, we hypothesized that *M. polymorpha* rhizoids may display similar alterations to those described for *A. thaliana* root hairs under Pi starvation. In order to test such hypotheses, we monitored rhizoid development at each time-point of the experiment. Our results showed that at 7 days after sowing (DAS), gemmae developed into young thalli in both +Pi and −Pi conditions (*Figure 2a*). At this stage, we already observed differences in the length of rhizoids under −Pi when compared to +Pi (*Figure 2a*). The differences between +Pi and −Pi conditions were dramatic in both rhizoid length and density at 21 DAS (*Figure 2a*). In *A. thaliana*, upon Pi starvation, the increase in root hair density correlates with the excretion of Purple Acid Phosphatases (PAPs) to solubilize Pi from organic compounds (del Pozo et al., 1999). We explored if *M. polymorpha* rhizoids excrete PAPs by using the phosphatase’s substrate EFL-97 or *2-(2’-phosphoryloxyphenyl)-4(3H)-quinazolinone* (Huang et al., 1992), which is converted to *2-(2’-hydroxyphenyl)-4(3H)-quinazolinone*, a fluorescent product which emits a green signal at 530 nm wavelength. For this experiment, gemmae were grown under −Pi and + Pi in solid media, both supplemented with 25 μM of ELF-97 (*see methods for details*). After transference to media and poured into micro chambers to follow each analyzed gemmae, we recorded confocal sections at 0, 24, 48, 72, 120 and 168 hours after sown (HAS). Our results are shown in Figure 2b, where the green signal corresponds to PAP enzymatic activity on ELF-97, while the red signal corresponds to chlorophyll autofluorescence. At 0 HAS no evident green signal was observed among treatments. By 24 HAS, a discrete green signal was observed in both +P and −Pi conditions. However, at 48 HAS a stronger green signal was observed in gemmae grown in −Pi, not only in rhizoid cells but also in surrounding cells. Such stronger signal was not only observed in rhizoid cells but in surrounding cells that show chlorophyll autofluorescence in red (*Figure 2b magnifications*). At this time point, +Pi grown gemmae also showed green signal in the rhizoids, however, the green fluorescence intensity in these cells was lower and no evident green signal colocalizing with red signal was observed (*Figure 2b magnifications*). The patter of the green signal was even more intense at 72 and 120 HAS under −Pi conditions, when compared to those observed in gemmae grown under +Pi conditions at the same time points (*Figure 2b magnifications*). We also observed in −Pi grown plants that several chlorophyll-producing cells, in the region of the thallus, where rhizoids are formed, also showed a more intense green signal when compared to the control (*Figure 2b magnifications*). Our phenotypic analysis revealed that the proper development and growth of plants, grown under low Pi conditions, was also drastically altered when compared to those of plants grown in +Pi (*Figure 2a*). The biomass production was affected lightly after 7 DAS and was clearly lower at 14 and 21 DAS under −Pi conditions relative to the control (*Figure 2c*). In addition, we observed an evident change in the green color characteristic of the thallus to a strong red/purple color, suggesting that accumulation of phenylpropanoids (*e.g*. aunoridins) in response to low Pi conditions is a conserved feature of land plants in response to scarce Pi environments (*Figure 2a*). The previously described phenotypes correlated with a significant decrease in free Pi in plants grown under low Pi conditions, in comparison with the control (*Figure 2d*). Collectively, our phenotypic analyses showed that Pi deficiency causes alterations in early thallus development, some of which were analogous to those previously described for *A. thaliana* and other land plant species, suggesting that several phenotypic and physiological responses triggered by this stress are conserved across streptophytes.

**Figure 2.**
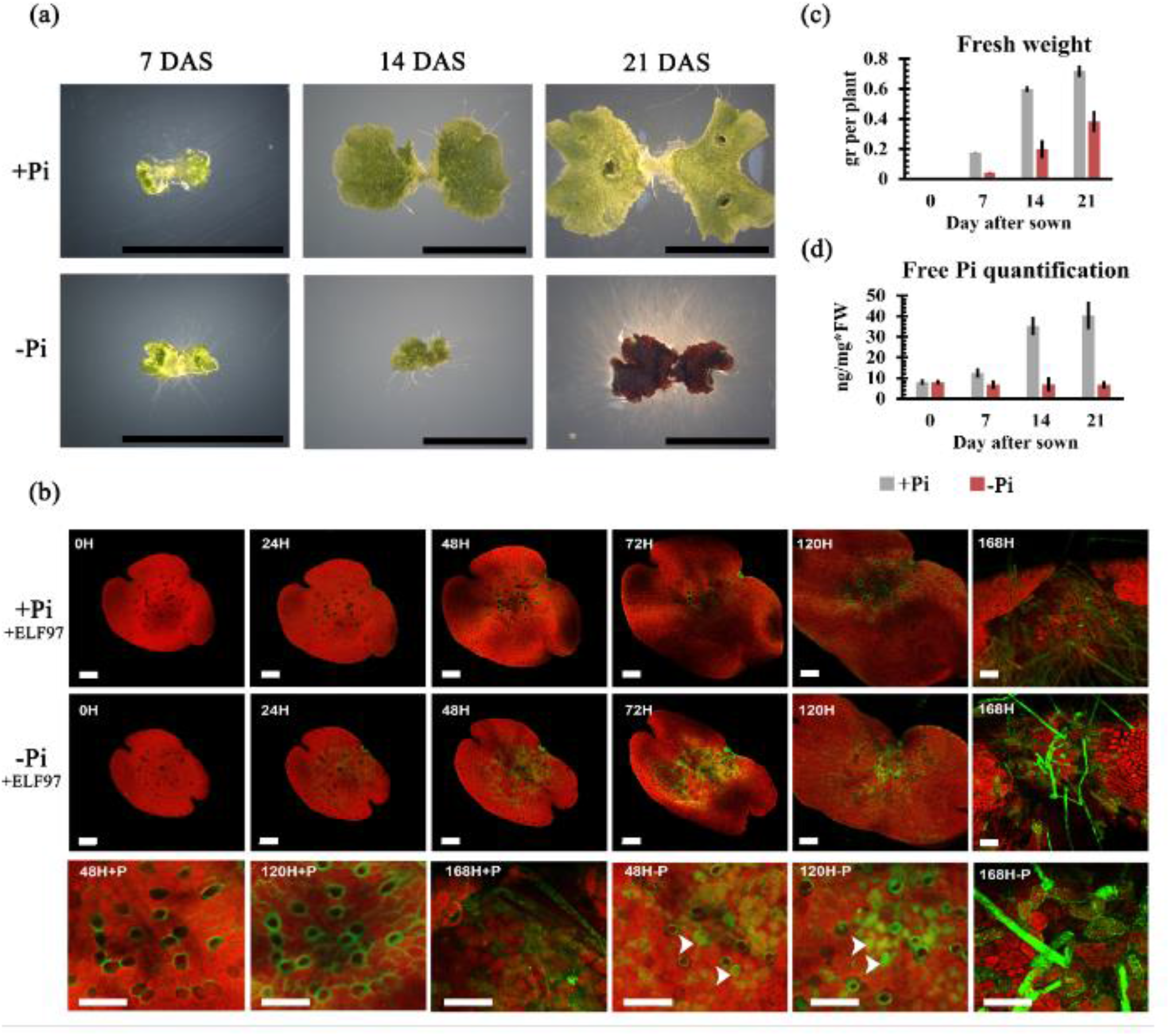
Phenotypical and physiological effect of phosphate availability in *Marchantia polymorpha*. (a) Morphological effect of +Pi (500 μM) and −Pi (10 μM) status on talli development in plants to 7, 14 and 21 DAS grown under continuous stressful conditions (Black lines = 5mm). (b) *In vivo* analysis of acid phosphatase activity with ELF-97 in gemmae after 0, 24, 48, 72 and 120 hours after sowing. (c) Fresh weight (N = 30 plants) and (d) Free Pi quantification of plants exposed to high and low Pi conditions after 7, 14 and 21 DAS (N = 30 plants). All bars in confocal images, panel (b) =100um.

### 4) Transcriptional dynamics in response to Pi availability

The transcriptional response upon Pi starvation has been characterized in diverse angiosperms, such as *A. thaliana, O. sativa, Z. maize* and other angiosperms, where the transcriptional responses induced by low Pi availability are broadly conserved (Misson et al., 2005; Secco et al., 2013; Calderón-Vázquez et al., 2011; Zeng et al., 2016). However, the transcriptional response to low Pi has not been explored in earlier diverging lineages of land plants, such as liverworts. To gain insight on the transcriptional program of young plants relative to Pi availability, we employed an RNA-seq strategy in thalli derived from gemmae 10 DAS (*Figure 3a*). Plants were grown previously on Pi-sufficient liquid media for 10 days and subsequently transferred to either high (500 μM) or low Pi (10 μM) MS liquid media (*Figure 3a*). We defined the times for tissue collection and RNA isolation at 12, 24 and 150 hours post transference (HPT) (*Figure 3a*), based on the following criteria: 1) The induction of phosphatase activity in low Pi conditions observed at 24 h after transfer, suggesting early transcriptional activation (*Figure 2b*), 2) first distinct phenotypic changes, *e.g*. rhizoid phosphatase activity, were observed at 168 h (7 d) after sowing (*Figure 2b*), 3) the morphological changes in the number and length of rhizoid cells were qualitative evident at 168 h (7 d) after sowing (*Figure 2a and Supplementary Figure S25*), and 4) the acidification of media, probably by the exudation of organic acids and proton extrusion was observed at 7 DAS in Pi limited conditions (*Supplementary Figure S26*). In addition, biological replicates were obtained in an independent experiment under the same growth conditions and RNA collection. Our transcriptome resulted in a total of 274,795,583 reads, after removing poor quality reads and adapters. To estimate the gene expression profile, we mapped the reads using the DNA subway platform from cyverse (https://dnasubway.cyverse.org/) against the reference genome V3.1 (Bowman et al., 2017). To determine differentially expressed genes (DEGs) we contrasted libraries from the low Pi conditions against those from plants grown under high Pi conditions. We applied a fold change of [-1.5 < FC > +1.5] and a q-value < 0.001 as a selection criteria. From a total of 19,138 genes, 6,742 (35.22%) were differentially expressed (green bar *in figure 3b*), while 12,396 genes did not show significant differential expression (grey bar in *Figure 3b*). Then, we classified the total DEGs in up or down-regulated at each specific time point (12, 24 and 150 HPT). Bar plot in Figure 3c showed in orange 1,463; 1,196 and 2,258 up-regulated genes. While in blue we found 1,202; 1,604 and 2,570 down regulated genes at 12, 24 and 150 HPT, respectively. In order to define time-specific and shared DEGs, we performed a Venn diagram analysis for up (orange) and down-regulated genes (blue) (*Figure 3d*). The up-regulated time-specific DEGs were 599, 625 and 1,639 genes at 12, 24 and 150 HPT and for down-regulated 490, 725 and 1825 at 12, 24 and 150 HPT, respectively (*Figure 3d*). Also, 152 up-regulated genes were shared by the three time points, while 258 down-regulated genes were shared among all times. While the remaining DEGS were shared between two timepoints. For example, in the up-regulated group, samples from 12 and 24 HPT shared 332 DEGs, 24-150 HPT shared 87 and 12-150 HPT shared 380 genes. On the other hand, the down regulated group showed 289 DEGs shared between 12 and 24 HPT, 322 between 24 and 150 HPT and 165 DEGs shared among 12 and 150 HPT (*Figure 3d*). In order to reveal the putative functions of DEGs in response to Pi deprivation, we performed a Gene Ontology (GO) enrichment analysis. The total universe of up regulated DEGs was used to define enriched subcategories from the main category “*Biological Processes*” (*BP*) with a threshold of *p*-value < 0.001. Then we used the ClueGO software (Bindea et al., 2009) to cluster the over-represented categories in a network, where the circle size correlates to p-value and grey lines define interactions between subcategories (*Figure 3e*). The major enriched category is the *Aromatic acid family metabolic process*, it forms a subnetwork including subcategories such as *Indole-containing compound metabolic process*, *Aromatic amino acid family biosynthetic process*, *Cellular amino acid catabolic process*, *Organic acid metabolic process* and others (*Figure 3e*). Into these subcategories we found genes associated with auxin biosynthesis, organic acid cycle and exudation. Other subnetwork highly enriched includes *Polysaccharide catabolic process, Starch catabolic process, Glucan catabolic process* and others. Interestingly, the *Ion transport* category formeds another subnetwork which includes *Phosphate ion transport*, *Iron transport* and other subcategories (*Figure 3e*). In these subcategories, we found genes involved in Pi and Fe transport, and putative Pi transporter genes such as homologues of *PTB* and *PHT* were induced in response to low Pi availability. Another category was the *Flavonoid biosynthesis* and *Chalcone biosynthetic process*, where genes related to auronidin biosynthesis were found (Albert et al., 2018). The transcriptional activation of these genes correlates with the low Pi phenotype observed (dark purple color on thallus *Figure 2a*). In addition, we found conservation of some of the typical responses to low Pi availability such as the induction of genes underlying the plant hormone genetic networks, organic acid exudation, iron transport, Pi transport and cellular responses to Pi starvation (*Figure 3e*). We then searched for enriched functional categories of up regulated genes specific for each time point, these results are available in the *supplementary Figures S27, S28 and S29*. The analysis was performed using a similar procedure to the global GO analysis, but the threshold was adjusted to *p-value* < 0.05. Our results for the 599 DEGs specific at 12 HPT (*Figure 3d*) showed enrichment in 66 functional categories which included *Response to hormones, Auxin polar transport, Response to cytokinin, Auxin-activated signaling pathway* and *Response to jasmonic acid (Supplementary Figure S27*). Other interesting categories were *Carboxylic acid biosynthetic process*, *Cellular response to superoxide*, *Chalcone biosynthetic process*, *Regulation of cell morphogenesis involved in differentiation* and *Cellular response to phosphate starvation (Supplementary Figure S27*). We found 570 diverse functional categories enriched in the 725 DEGs specific for 24 HPT (*Figure 3d*). Some of these subcategories related to plant hormones such as *Response to abscisic acid, Response to ethylene* and *Response to salicylic acid (Supplementary Figure S28*). Also, we found categories related to processes which have been previously associated with the Pi-local response including *Meristem development, Defense response by callose deposition in cell wall* and *Root system development (Supplementary Figure S28*). Other interesting functional categories found were *Response to lipid, Iron ion homeostasis* and *Sulfate assimilation*, the rest of the varied subcategories are represented in (*Supplementary Figure S28*). Our analysis of the 639 specific DEGS of the 150 HPT time point revealed enrichment in 250 functional subcategories (*Supplementary Figure S29*). Some of the most interesting subcategories were *Carboxylic acid transport*, *Citrate metabolic process*, *Programmed cell death* and *Response to aluminum ion (Supplementary Figure S29*). We also found *Organophosphate ester transport*, *Lipid modification*, *Lipid oxidation* and *Inositol metabolic process (Supplementary Figure S29*). Together, our time-specific GO analysis reveals that the transcriptional response of *M. polymorpha* to cope with Pi starvation is dynamic an involves the upregulation of genes associated to several key processes, some of which were time-specific and conserved partially with those described in *A. thaliana, O. sativa* and *Z. mays* (Misson et al., 2005; Secco et al., 2013; Calderón-Vázquez et al., 2011; Zeng et al., 2016). This data suggests that our transcriptome captures the response of at early (12 and 24 HPT) and late (150 HPT) time points upon Pi limitation in *M. polymorpha*. This contributes to understanding the transcriptional dynamics in function of Pi availability and shows a partial conservation with previous characterizations in other flowering plants.

**Figure 3.**
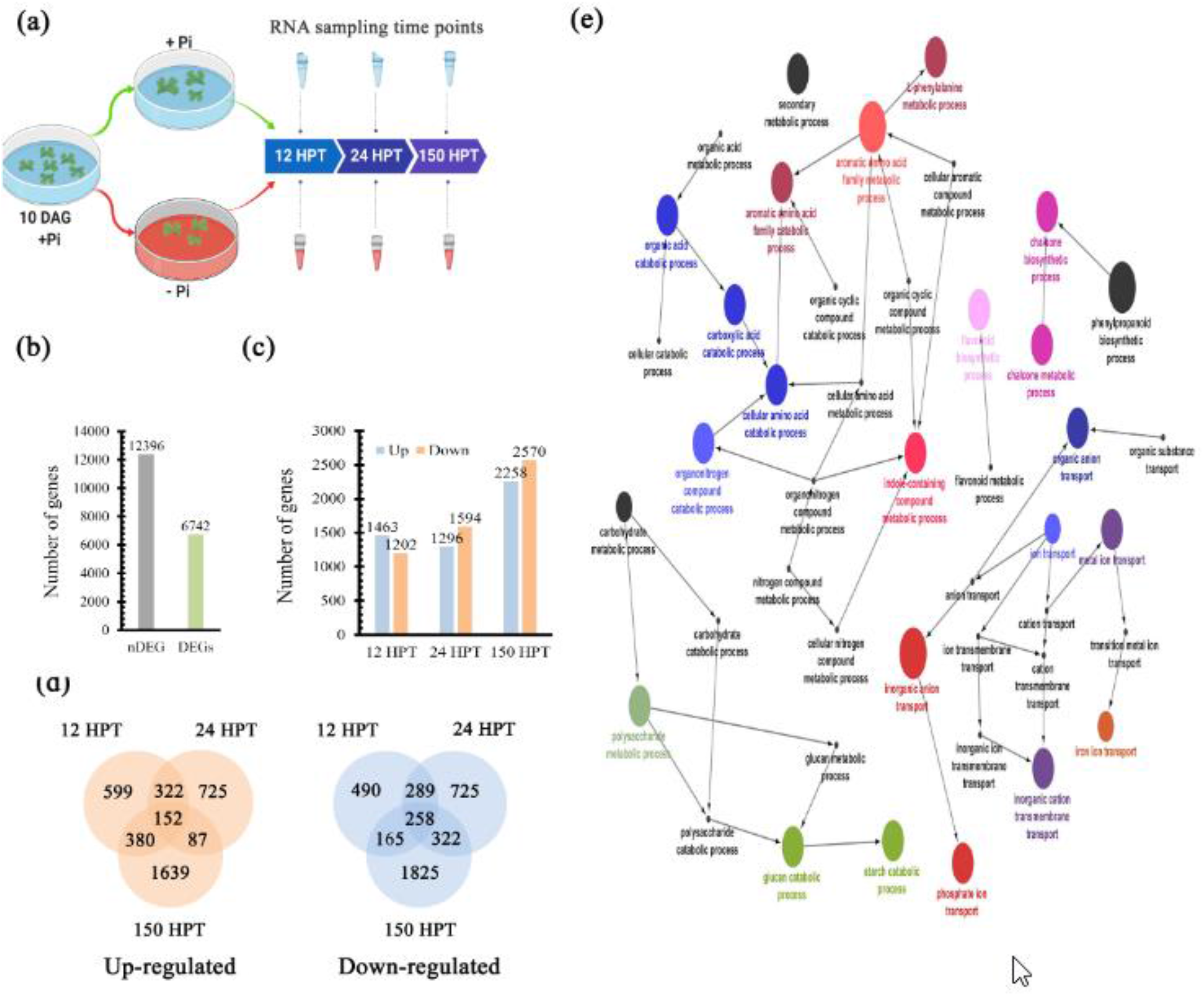
Dynamics of transcriptional responses to Pi starvation. (a) Graphic description of the experimental design for the RNAseq approach. (b) Differentially expressed genes (DEGs) and non-differentially expressed genes (nDEGs). (c) Bar plot showed the up-regulated genes in light red and the down-regulated ones in green for each time of transcriptome. (d) Venn diagrams for up-regulated (Red) and down-regulated (green) genes for each condition. (e) Clustering of over-represented gene ontology (GO) categories in the genes up-regulated in response to low Pi availability, color clustered similar categories and circle size correlates with the P-value of enrichment test.

### 5) Transcriptional regulation of the local and systemic Pi-sensing modules

In order to explore how Pi sensing occurs in *M. polymorpha*, we analyzed the transcriptional behavior of genes involved in local and systemic perception networks. Interestingly, only Mp*PHR1* was up-regulated in low Pi, in contrast to the other two MYB-CC genes (*Supplementary Figure S30*). This suggests that Mp*PHR1* is functionally equivalent to *AtPHR1*. Meanwhile, Mp*SPX* was up-regulated across all time points in response to low Pi similar to that observed for AtSPX1/2. Respect to the local perception network, we found that Mp*STOP2* was significantly down-regulated at 12 HPT and up-regulated at 24 and 150 HPT, under Pi deficiency. Based on its transcriptional activation, we hypothesized that Mp*STOP2* (Mapoly0083s0069) may be functionally equivalent to *AtSTOP1*. Mp*ALMT3* (Mapoly0100s0061) and Mp*ALMT2* (Mapoly0070s0016) were significatively down regulated at 12, 24 and 150 HPT, even when Mp*STOP2* was up-regulated (*Supplementary Figure S30*). Although we identified 13 loci encoding multicopper oxidases (Mp*LPR1* to Mp*LPR13*), only two of them, Mp*LPR1* (Mapoly0008s0168) and Mp*LPR3* (Mapoly0008s0270), were up-regulated in all three time points sampled in this work (*Supplementary Figure S30*). While Mp*LPR4* (Mapoly0030s0116) and Mp*LPR7* (Mapoly0186s0019) were up-regulated at 12 HPT and then down-regulated at 24 and 150 HPT. Mp*LPR11* (Mapoly0217s0007) was up-regulated at 12 and 24 HPT, then down-regulated at 150 HPT (*Supplementary Figure S30*). Only Mp*LPR2* (Mapoly0004s0015) was down-regulated in the three time points analyzed (*Supplementary Figure S30*). On the other hand, the CLE peptide family in *M. polymorpha* contains two members (Mp*CLE1* and Mp*CLE2*) (Hirakawa et. al., 2019), under low Pi conditions Mp*CLE1* (Mapoly1011s0001) was down-regulated at 12 and 24 HPT, while Mp*CLE2* (Mapoly0084s0052) was up-regulated at 12, 24 and 150 HPT (*Supplementary Figure S30*). Taken together these results indicate that despite the differences, at morphological and genetic levels, with Arabidopsis, *M. polymorpha* displays a partially conserved transcriptional response of both, local and systemic, Pi sensing pathways.

### 6) Low Pi promotes the induction of genes related to organic acid synthesis and exudation

To unveil if genes involved in organic acid synthesis and exudation were affected in response to low Pi conditions, the transcriptional profiles of key genes such as the malate synthase (MLS), citrate synthase (CIS), malate transporters (ALMT) and citrate transporters (MATE) were analyzed (*Supplementary Figure S31*). Our search in the *M. polymorpha* genome with the Pfam identifiers PF00285 for citrate synthase, PF01274 for Malate synthase and PF01554 for the MATE transporter, revealed five loci encoding putative *MALATE SYNTHASE (MpMLS1 to MpMLS5*), from which the transcripts of Mp*MLS2* (Mapoly0016s0095), Mp*MLS3* (Mapoly0023s0108) and Mp*MLS5* (Mapoly0154s0035) were down regulated under −Pi conditions (*Supplementary Figure S31*). On the other hand, we found 9 loci annotated as *CITRATE SYNTHASE* (Mp*CIS1 to* Mp*CIS9*), four of them were differentially expressed under low Pi conditions. Mp*CIS1* (Mapoly0063s0010) was upregulated at 12 HPT and downregulated at 24 and 150 HPT, while Mp*CIS2* (Mapoly0071s0012) and Mp*CIS5* (Mapoly0111s0039) were down-regulated at 12 and 24 HPT, while up regulated at 150 HPT (*Supplementary Figure S31*) and Mp*CIS6* (Mapoly0200s0007) was down-regulated at the three times analyzed (*Supplementary Figure S31*).

Regarding malate transporters, Mp*ALMT3* and Mp*ALMT4* were down-regulated, as previously mentioned. This observation correlates with reduced levels of expression of malate synthesis enzymes and suggests a reduced production and exudation of malate under low Pi conditions. In the case of genes related citrate transport, we identified 15 loci encoding putative MATE-type transporters (MpMATE1 to MpMATE15), (*Supplementary Figure S31*). In Arabidopsis some members of this family of transporters are involved in citrate exudation (Liu et al., 2009). Interestingly, Mp*MATE1* (Mapoly0002s0036), Mp*MATE3* (Mapoly0005s0170) and Mp*MATE10* (Mapoly0041s0128) were upregulated in all time points sampled (*Supplementary Figure S31*). Additionally, other six members of the MATE family were up-regulated in at least one time point sampled (*Supplementary Figure S31*). These results suggest that *M. polymorpha* promotes preferentially the synthesis and efflux of citrate rather than malate to cope with low Pi.

### 7) Pi starvation induces the expression of genes involved in lipid turnover

To explore if phospholipid remodeling takes place in *M. polymorpha* in response to Pi limited conditions as previously reported for *A. thaliana* (Essigmann et al., 1998; Andersson et al., 2003; Cruz-Ramirez et al., 2006), we searched for putative orthologs of key enzymes involved in phospholipid degradation and glycolipid synthesis (*Figure 1a*). Our results revealed 7 loci encoding putative orthologs of type D phospholipases (PLDs), our phylogenetic reconstruction showed that *M. polymorpha* PLDs formed a basal clade with PLD sequences of bryophytes (*Supplementary Figure S13*). Among the *M. polymorpha* PLD-encoding loci, we found that Mp*PLD1* (Mapoly0191s0011) is up regulated at the three time points analyzed and displayed the strongest FC value at 150 HPT (*Supplementary Figure S32*). This suggests that Mp*PLD1* is probably the functional equivalent of *AtPLDz2*, which is also highly up-regulated in response to Pi-deprivation (Cruz-Ramirez et al., 2006). Although other members of the *PLD* family genes in *M. polymorpha* were down and up-regulated at different time points, showed lower FC values than those of Mp*PLD1* (*Supplementary Figure S32*). In the case of enzymes involved in galacto and sulpholipid synthesis, we found single genes encoding them, Mp*PAH*, Mp*DGDG*, Mp*DAG*, Mp*MGDG* and Mp*SQD*. Among them, Mp*DGDG* and Mp*MGDG* did not show significant changes in their expression relative to the availability of Pi, while Mp*DAG* was down-regulated at 12 and 24 HPT and up-regulated at 150 HPT. In contrast, Mp*SQD* was up-regulated at 12 and 24 HPT and down-regulated at 150 HPT in low Pi (*Supplementary Figure S32*). Altogether, this data show that a group of genes, which homologs in Arabidopsis are involved in lipid turnover, are up-regulated in response to Pi availability in *M. polymorpha*.

### 8) Transcriptional behavior of genes involved in ROS synthesis in Pi starved thallus

Previous studies have associated reactive oxygen species (ROS), such as hydroxyl radical (OH^-^), superoxide (O^-^) and hydrogen peroxide (H_2_O_2_), to the regulation of plant development by acting as secondary messengers (Mitter et al., 2011). It has been shown, in Arabidopsis, that NADPH OXIDASE (NOX1) or *RESPIRATORY BURST OXIDASE HOMOLOG* (*AtRBOH*), a major enzyme in the production of ROS, is required for the appropriate root hair developmental patterning (Foreman et al., 2003; Carol et al., 2005). Moreover, transcriptional studies of *A. thaliana* upon Pi starvation revealed that peroxidase-encoding genes were significantly up-regulated (Mora-Macias et al., 2017). In order to explore the conservation of this response in the early divergent land plant *M. polymorpha* in response to Pi starvation, the transcriptional profiles of *NOX/RBOH* and peroxidase-encoding genes were analyzed. The Pfam identifiers PF08414 (NADPH oxidase) and PF00141 (Peroxidase) were used to identify the putative *M. polymorpha* orthologs. We found two loci for *NADPH OXIDASE* (Mp*NOX1* and Mp*NOX2*) (*Supplementary Figure S34*). The expression of Mp*NOX1* (Mapoly0258s0001) was up-regulated at 12 and 150 HPT, but down-regulated at 24 HPT (*Supplementary Figure S34*), while Mp*NOX2* (Mapoly0046s0097) was down-regulated at the three time points sampled (*Supplementary Figure S34*) In contrast, we found 221 loci putatively encoding peroxidases (PRX), from which 100 loci showed differential expression in our transcriptomic approach (*Supplementary Figure S33*). Transcripts of 24 loci were constantly down-regulated and 14 up regulated in the three time points analyzed, while the remaining 62 were up or down-regulated in at least in one of the time points sampled (*Supplementary Figure S33*). The transcriptional induction of Mp*NOX1* under low Pi conditions suggests that *M. polymorpha* promotes the generation of superoxide radical (O^-^). Also, the strong induction of several Mp*PRX* genes might correlate with ROS production under low Pi conditions and could be involved in the changes observed in rhizoid development upon Pi scarcity.

### 9) Validation of transcriptome by qPCR

In order to evaluate the quality of our transcriptome, 7 genes were selected for qPCR analysis. The putative orthologs of the selected genes have been associated with relevant aspects of Pi starvation responses in diverse plant species, such as Pi sensing, cellular response to Pi, organic acid biosynthesis and transport, actin was used as control (*Figure 4a*). For our qPCR assays we used cDNA derived from total RNA sampled at the same time points and growth conditions used for our transcriptome. The transcriptomic profiles of the selected genes are shown in Figure 4a, up-regulated genes are depicted in color orange down regulated ones in blue and unchanged ones in black. For the internal Pi perception, our transcriptome showed that Mp*PHR1* and Mp*SPX*, were up-regulated in the three time points analysed under low Pi conditions (*Figure 4a*). In contrast, our qPCR analysis showed that Mp*SPX* transcripts under low Pi conditions were less abundant at 12 HPT, but increased drastically at 24 HPT and 150 HPT, compared to high Pi conditions as the control (*Figure 4b*). In the case of Mp*PHR1*, transcript levels increased significatively in response to Pi starvation at the three time points tested (*Figure 4e*). These qPCR results denote a high coherence with our transcriptomic data for these genes. Regarding organic acid biogenesis, our transcriptomic approach showed that transcript levels of Mp*MLS3* were downregulated at all timepoints sampled, while those of Mp*CIS1* were down-regulated at early times (12 and 24 HPT) and up-regulated at 150 HPT (*Figure 4a*). The qPCR analysis revealed that transcript levels of Mp*MLS3* were significantly lower in low Pi at 12 and 150 HPT respect to the control, while no significant differences between high and low Pi conditions were observed at 24 HPT (*Figure 4c*). Meanwhile, qPCR results for Mp*CIS1* showed no significant differences at early times (12 and 24 HPT), but they highly increased at 150 HPT under low Pi conditions (*Figure 4d*). Finally, we analyzed the transcriptional behavior of genes whose putative orthologs in *A. thaliana* are involved in the local perception of Pi availability. Our transcriptome showed that Mp*CLE1* was down-regulated at early times (12 and 24 HPT) whereas the Mp*CLE2* was upregulated in the three points sampled, in response to low Pi (*Figure 4a*). The qPCR data for Mp*CLE1*, showed low transcript abundance in −Pi conditions at the three time points tested (*Figure 4f*), while transcript levels for Mp*CLE2* were high under low Pi conditions at the three time sampled (*Figure 4g*), showing a high correlation with the expression pattern observed for this gene in our transcriptome approach. In the case of Mp*LPR1*, the transcriptome data showed that transcription of this gene is not altered between treatments at early times (12 and 24 HPT) and is up-regulated under low Pi at 150 HPT (*Figure 4a*). Our qPCR analysis corroborated this expression pattern, only after 150 HPT Mp*LPR1* showed high transcript levels under low Pi conditions (*Figure 4h*).

**Figure 4.**
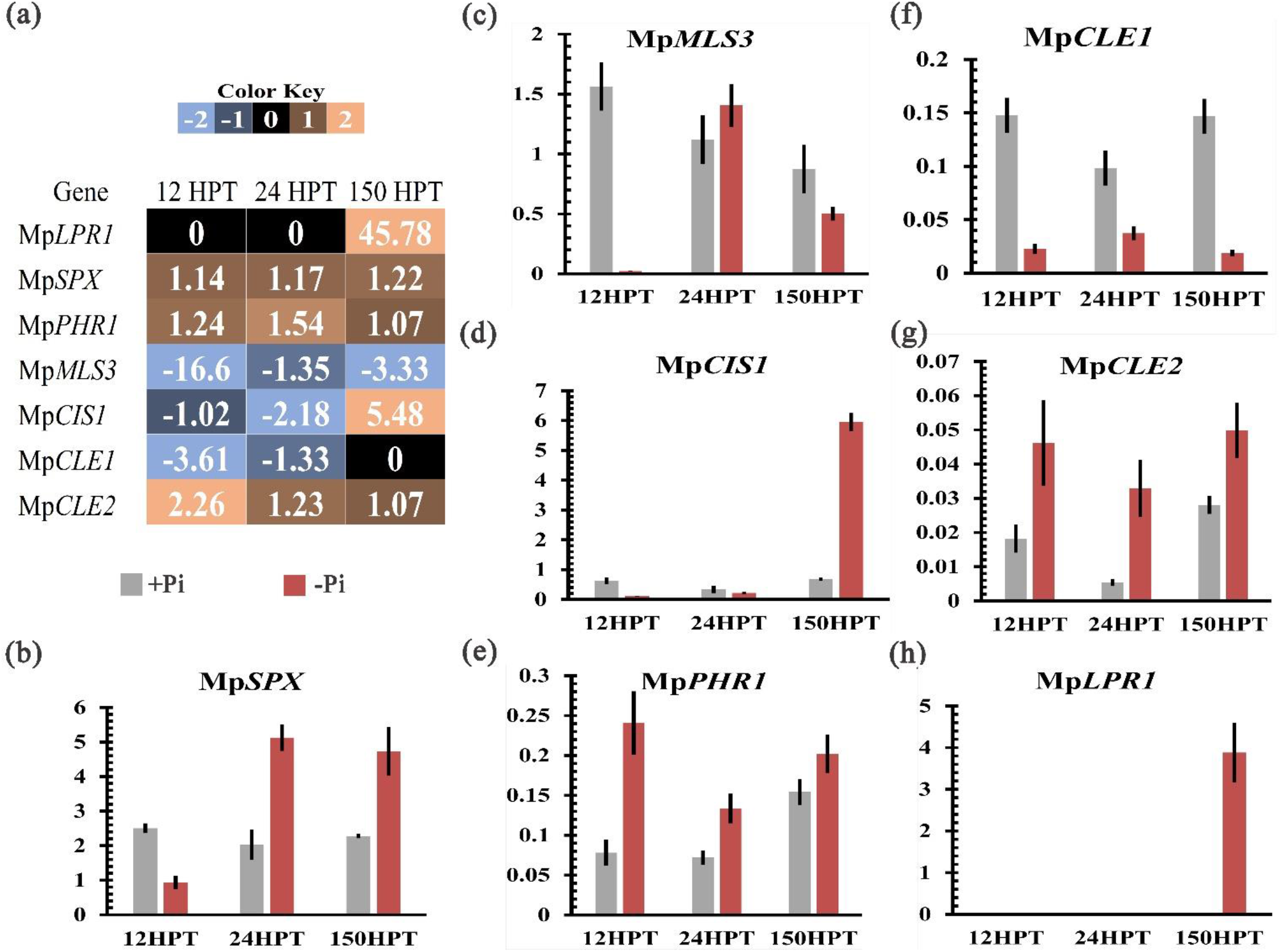
Validation of transcriptome by Quantitative real-time qPCR. (a) Heat map showed the transcriptional profile of Mp*SPX*, Mp*PHR1*, Mp*CLE2*, Mp*LPR1*, Mp*CIS1*, Mp*CLE1* and Mp*MLS3* resulted from our transcriptome. Gradient of colors represents light blue down-regulated, black means no change in expression and orange represents up-regulation. (b-h) Transcript levels of Mp*SPX*, Mp*PHR1*, Mp*CLE2*, Mp*LPR1*, Mp*CIS1*, Mp*CLE1* and Mp*MLS* genes at 12, 24 and 150 HPT, determined by qRT-PCR. The Y axis for all bar plots represent relative expression.

Overall, our qPCR analysis validated the transcriptional behavior of most of the genes selected, indicating that our transcriptome approach is robust and that the reported expression patterns indeed reflect the response of *M. polymorpha* to low Pi.

### 10) Determination of PHR1 Binding Sites (P1BS) enrichment on DEGs

Previous studies in *A. thaliana* reported the enrichment of the P1BS *cis* regulatory element (CREs) in the promoter regions of *AtPHR1* target genes (Thibaud et al., 2010; Bustos et al., 2010). Such genes were transcriptionally impaired in the loss of function mutant *phr1* and the double mutant *phrphl1*, unveiling the relevance of the P1BS in the control of Pi starvation responses (Bustos et al., 2010).

We hypothesized that some of the genes which are differentially expressed in response to Pi starvation in *M. polymorpha*, may correlate with the presence of the P1BS motif in their promoter regions. In order to test this idea, we surveyed for the presence and enrichment of the P1BS motif within a 3 Kb region upstream the transcription start site of DEGs and non differentially expressed genes (nDEGs).

First, we searched the P1BS consensus sequence (*GNATATNC*) in the promoters of the 6746 DEGs using HOMER (detailed in Methods) and found that 3259 of them harboured at least 1 P1BS. We then performed a motif enrichment analysis in the promoters of those 3259 DEGs and in the 10894 nDEGs using the Analysis of Motif Enrichment (AME, Version 5.1.1) (McLeay and Bailey, 2010) tool from the MEME suite (Bailey et al., 2009). AME identifies if the motif, in this case P1BS, is enriched in a set of sequences considered as primary (DEG and nDEG promoters) compared to a control set of sequences (the 16240 *M. polymorpha* annotated promoter regions). AME scores every sequence in terms of presence of P1BS and performs a statistical analysis to determine if the primary sequences have better scores than the control sequences. Our results show that 2429 promoters of DEGs are enriched for P1BS, while nDEGs for 946. Promoters from DEGs were then classified in up-regulated and down-regulated. From the total of up regulated DEGs (5499 promoters), 3531 have no P1BS, 1188 bear 2 P1BS and 780 have 1 P1BS (*Figure 5a*). We then explored the GO categories on the up-regulated transcripts which promoters have 1 and 2 P1BS, and found several subcategories highly associated with Pi starvation responses (PSR) such as *inorganic phosphosphate transmembrane transporter activity, triglyceride lipase activity, endoribonuclease activity and others* colored in red in the network (*Figure 5b*). While the subcategories not related directly to PSR appear colored in black (*Figure 5b*). In the case of down-regulated DEGs, we analysed a total of 1247 promoter regions for enrichment in the P1BS CRE and found that 142 have 1 P1BS, 319 have 2 P1BS and 786 have no P1BS (Figure 5c). Respective GO analysis for those which contain the P1BS motif revealed subcategories that are linked to the pathogen response, *i.e. endopeptidase inhibitor type C*, in blue (Castrillo et al., 2017), and other subcategories are colored in black (*Figure 5d*). The last group contained genes whose expression did not show significant changes (10894 nDEGs), and we found that only 500 and 546 promoters have 1 and 2 P1BS, respectively (*Figure 5e*), while the remaining 9803 have no P1BS. The GO analysis of this group revealed categories that in Arabidopsis have been related to Pi starvation (in red) and others stress responses which have been previously related to AtPHR1, i.e. *Zinc ion binding and inositol monophosphate phosphatase activity* (Briat et al., 2015; Li and Lan, 2015) (*Figure 5f*).

**Figure 5.**
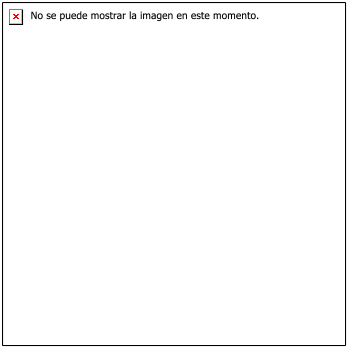
Genome-wide identification of PHR1 binding sites (P1BS) distribution on *M. polymorpha*. a) Pie chart showing the number of promoter regions of up-regulated DEGs which lack the P1BS motif (in gray) and those whose promoter has 1 (green) or 2 (blue) motifs, the same color code applies for those pie charts y c) and e). b) Illustrates the GO analysis of the up-regulated DEGs which contain the P1BS CREs, red color indicates the subcategories that are strongly associated to Pi starvation responses and in black those terms which have no evident link to the PSR. c) Pie chart showing the number of promoter regions of down-regulated DEGs which lack the P1BS motif, and those whose promoter has 1 or 2 motifs. d) The GO network for the down regulated genes with P1BS motifs, blue nodes indicate subcategories which have been related to PHR1 function in regulating genes involved in other stress conditions described in *A. thaliana* (Castrillo et al., 2017), dark nodes show categories with no evident link to the Pi starvation response. e) Pie chart showing the number of promoter regions of non-DEGs which lack the P1BS motif and those whose promoter has 1 or 2 motifs. f) The GO networks found after analysis the genes with P1BS motives in the nDEGs group, subcategories related to low Pi availability appear in red, whereas in black the subcategories non-associated to the Pi limitation response.

Overall these results show that the number of promoters enriched for P1BS is higher in up-regulated DEGs than in nDEGs. However, the upregulated and down-regulated promoter groups displayed similar tendencies for P1BS enrichment, but the GO analysis revealed that within the up-regulated group we mainly found subcategories related to PSR. By contrast, in the down-regulated targets we found overrepresentation of subcategories related to the response to pathogens. Hence, the presence of P1BS in the down-regulated and nDEGs suggests that MpPHR1 could be involved in the transcriptional regulation of diverse targets involved in a broad range of stress conditions (Briat et al., 2015; Li and Lan, 2015; Castrillo et al., 2017).

## DISCUSSION

### 1) Morpho-physiological responses upon Pi starvation

In the present work, the morphological and physiological responses of the rootless plant *M. polymorpha* to low Pi availability were described. Among these responses, we found an increase in the size and number of rhizoids under low Pi conditions (*Figure 6*). In *A. thaliana*, the increase in root hair density during Pi deprivation, has been described as a strategy to improve the assimilation zone in the rhizosphere (Peret et al., 2011). Hence, the modulation of rhizoid development under limited conditions of Pi could be involved in enhancing Pi uptake. It has been shown in *M. polymorpha* that rhizoid development is regulated by auxin signaling (Kato et al., 2015; Flores-Sandoval et al., 2015). Interestingly, our transcriptome showed that several genes related to auxin synthesis and signaling are differentially regulated in response to Pi starvation. It would be interesting to explore in future studies if the *M. polymorpha* enhanced rhizoid morphogenesis and growth phenotype observed during Pi deficiency invokes the action of auxin signaling genes, since auxin has been involved in root hair elongation in Arabidopsis (Lopez-Bucio et al., 2002) and rhizoid tip growth (Flores-Sandoval et al,., 2015).

**Figure 6.**
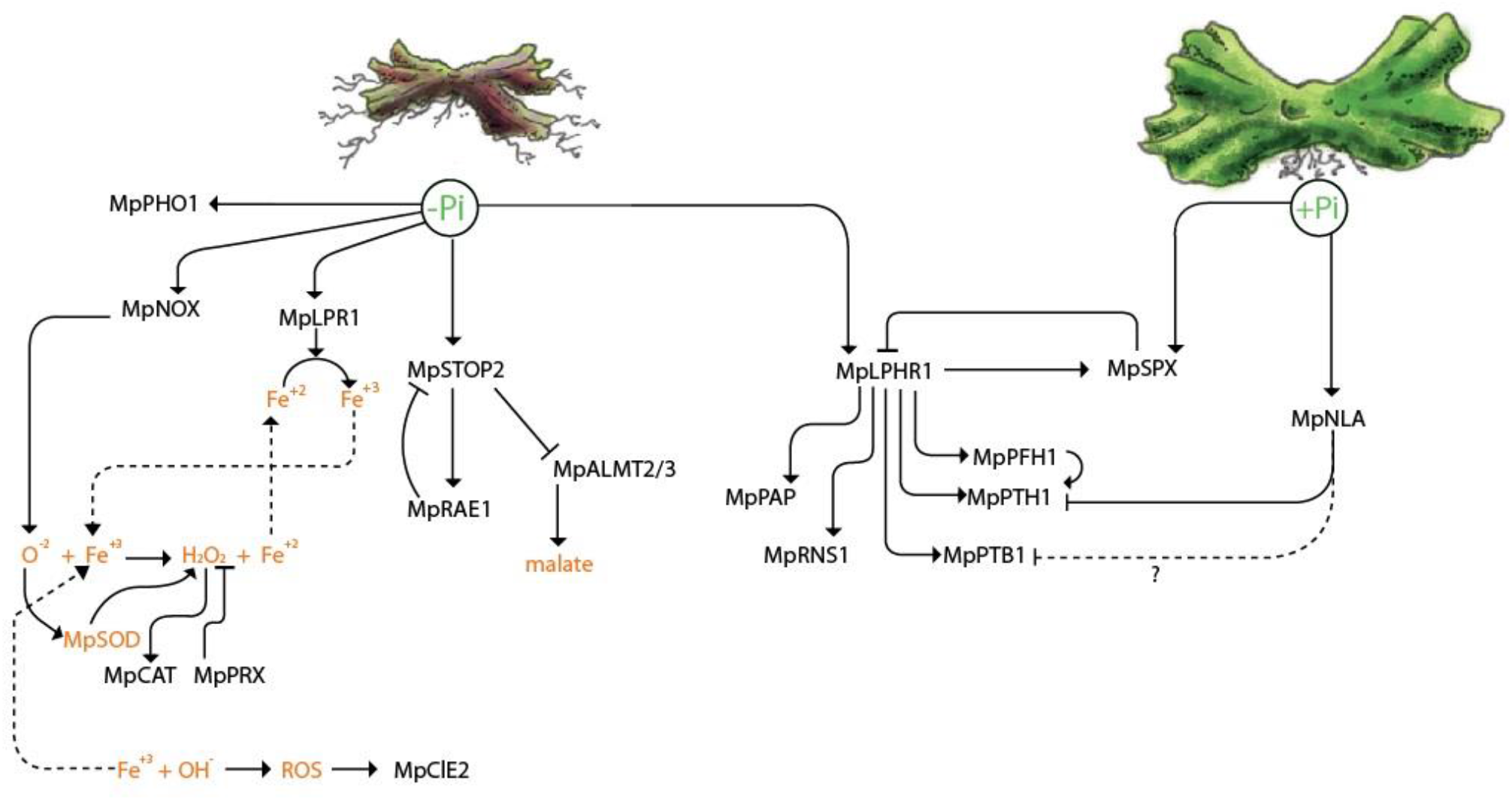
Model of the molecular response of *M. polymorpha* upon Pi starvation. Under low Pi conditions (left side of the figure) the expression of Mp*PHO1* is promoted. Mp*NOX1* induces the production of superoxide radical (O^-2^) which is catalyzed by SUPEROXIDE DISMUTASE (Mp*SOD*) into hydrogen peroxide (H_2_O_2_). Hydrogen peroxide is catalyzed by CATALASE (Mp*CAT*) into oxygen and water. Also, the peroxidases (Mp*PRX*) use hydrogen peroxide to carry out several reactions. On the other hand, the (O^-2^) plus Fe^+3^ carried out the Fenton-Weiss reaction to produce hydroxyl radical and Fe^+2^. This step coupled to the induction of Mp*LPR1*, a multicopper oxidase that leads to Fe^3+^ from Fe^+2^ under low Pi conditions, generates an oxidation-reduction process and probably impacts thallus development similar to what occurs in *A. thaliana* root. Here, we observed that the gene encoding the Mp*CLE2* peptide is up-regulated in specific times after growth in low Pi conditions, probably to modulate the maintenance and function of the meristem. Also, during low Pi deficiency, the expression of Mp*STOP2* is up-regulated, promoting the expression of its putative negative regulator the F-Box ubiquitin ligase Mp*RAE1*. Contrary to previous reports in vascular plants, Mp*ALMT2/3* are down-regulated under low Pi conditions. In agreement with this, we hypothesized that low Pi represses the expression of malate transporters to reduce the efflux on soil. On the other hand, the transcription factor Mp*PHR1* is induced under low Pi conditions and promotes the expression of MpPAPs and Mp*RNS1*, both genes involved in Pi scavenging. Mp*PHR1* promotes the expression of both Pi transporters MpPHTs and MpPTBs via the P1BS motives in their promoters. Additionally, we found that the Mp*PHF1*, an essential protein for the movement of MpPHT transporters from the ER to the PM, is under the control of Mp*PHR1*. In contrast, Mp*NLA is* expressed in high Pi conditions leading to the degradation of PHT and probably the degradation of the MpPTB transporters. Finally, among the Mp*PHR1* targets induced during Pi limitation we found Mp*SPX* which is a negative regulator of Mp*PHR1*. MpSPX is up-regulated under low Pi conditions under the control of MpPHR1 and another mechanism leads to its expression on high Pi availability.

We also found that biomass accumulation of thalli was negatively affected under low Pi conditions, as shown in *Figure 6*. This phenotypic change, in function of Pi availability, has been also observed in other plant models such as in *Arabidopsis thaliana, Oryza sativa and Zea mays* (Calderon-Vazquez et al., 2008; Lv et al., 2014; Liu et al., 2015). We also demonstrated in this study that free Pi levels are lower in plants grown under −Pi conditions compared to those grown in +Pi conditions, suggesting that the decrease of internal Pi amounts in *M. polymorpha* may trigger the expression of MpPHR1 targets, as reported in *A. thaliana* (*Figure 6*). This is supported by the fact that several putative homologs of this response such as Mp*PHR1* itself as well as Pi transporters from the Mp*PHT* and Mp*PTB* gene families are induced under low Pi conditions (Figure 6). In *A. thaliana*, the transcriptional activation of AtPHR1, under low Pi conditions, is controlled by AtARF7 and AtARF19 through three auxin response elements found on its promoter (Huang et al., 2018). Hence, it is possible that MpARFs could be involved in the transcriptional regulation of MpPHR1. However, future and detailed experiments are necessary to elucidate the putative upstream action of auxin, and other hormones, on the transcriptional regulation MpPHR1.

After 21 days of growth under low Pi conditions, thalli exhibited an intense red/purple color, which denotes accumulation of phenylpropanoids (*i.e*. auronidins in the case of *M. polymorpha*), a common phenotype associated to Pi deficiency across land plants (*Figure 6*). In *A. thaliana* the activity of key genes such as the *PHENYL AMONIOLIASE* (*AtPAL*) and *CHALCONE SYNTHASE* (*AtCHS*), both involved in anthocyanin biosynthesis, were up regulated under low Pi conditions (Misson et al., 2005). Our GO analysis on upregulated DEGs revealed enriched functional categories related to *Flavonoid biosynthesis* and *Chalcone metabolic processes*, both related to auronodin biosynthesis. The biochemical features of auronidin have been reported early, as well as the effects of uv light (Clayton et al., 2018) and nutrient deprivation (Albert et al., 2018; Kubo et al., 2018) in triggering its synthesis and accumulation (Berland et al., 2019). Here, we reported changes in the transcriptional patterns on key biosynthetic genes such as chalcone synthase, phenyl amonieliase and the transcription factor MpMYB14 (Mapoly0073s0038) mainly at 150 HPT (*Supplementary figure S35*). Our results show correlation between the up regulation of auronidin biosynthetic genes and the accumulation of this pigment in the thallus of plants grown in Pi deficiency.

Another important aspect of the Pi-starvation response reported in vascular plants, is the secretion of PAPs and OA to the rhizosphere. Our *in vivo* analysis of phosphatase activity revealed a strong induction of PAP’s activity in response to limited Pi, as previously reported for *Z. mays, O. sativa* and *C. reinhardtii* (Moseley and Grossman, 2009; González-Muñoz et al., 2015; Mehra et al., 2017). In the case of OA exudation, we observed media acidification occurring in thalli growing under low Pi conditions through an indirect measurement with bromocresol purple (*Supplementary Figure S26*), suggesting that OA efflux and proton extrusion is enhanced in response to Pi starvation. In agreement with this physiological evidence, the induction in the expression of genes encoding citrate synthase and MATE transportes was observed in our transcriptome (*Supplementary Figure S31*). In contrast, the malate synthase and ALMT transporters were down regulated (*Figure 6*). Collectively, our results suggest a partial conservation in *M. polymorpha* in the induction of OA production in response to low Pi. However, the transcriptional evidence suggests that, in order to cope with Pi scarcity, *M. polymorpha* preferentially promotes the synthesis and exudation of citrate rather than of malate, but further experiments are necessary to test this hypothesis.

### 2) Evolutionary perspective of Pi starvation responses along Viridiplantae

Pi sensing capacity is one of the major tools of plants to cope with Pi scarcity. In *A. thaliana* two independent networks involved in the perception of internal and external Pi levels have been described. Our evolutionary analysis unveiled a high conservation of the PHR1-SPX1 module for the internal Pi sensing networks in *M. polymorpha*, we found that putative orthologous genes are conserved since from chlorophyte algae (*Figure 1a; Figure 6*). This is a relevant finding, because it suggests an ancestral conservation in the negative regulatory mechanism of PHR1 by the SPX proteins (*Figure 6*). However, the functional conservation of the MpPHR1-MpSPX module, as a function of Pi availability, needs to be explored in future work (*Figure 6*). On the other hand, we found that the STOP1-ALMT1 regulatory network, which modulates the external Pi sensing, is a key innovation of streptophyte lineages (*Figure 1a; Figure 6*). We found that a putative STOP1 homolog is present from streptophyte algae, before the last common ancestor of land plants appeared. Therefore, we speculate that the STOP1-ALMT1 network was an innovation that influenced OA efflux and modified earth crust during the silicate weathering period, perhaps helping to reduce the levels of atmospheric CO2 (Reinhard et al., 2017).

The replacement of phospholipids by glycolipids in the lipidic bilayer of cell membranes is a well-described mechanism to deal with Pi scarcity which is highly conserved, even in bacteria (Alcaraz et al., 2008). We found a strong conservation of *XPL, PHA, DAG, MDG, DGDG, MGDG* and *SQD* putative orthologous genes from chlorophyte algae, streptophyte algae and *M. polymorpha*. However, genes encoding *PHOSPHOLIPASE alpha* (*PLA*) and *PHOSPHOLIPASE D* (*PLD*) were not found in the chlorophyte genomes analyzed. We hypothesized that other phospholipases, such as *PHOSPHOLIPASE C* (*PLC-L*) and *PHOSPHOLIPASE B* (*PLB*), could be converting phospholipids into DAG for the production of sulfo and galactolipids, as it was reported for *C. reinhardtii* under nitrogen limitation (Gargouri et al., 2015). Other key genes which we found highly conserved are the putative PHT1 transporters, found along plant phylogeny from chlorophyte algae to vascular plants (*Figure 1a*). However, we found that sodium dependent Pi transporters (PTB) are present only in green algae and *M. polymorpha*, suggesting that vascular plants lost this type of transporters. In agreement with these findings, the conservation of PTB transporters in *M. polymorpha* was reported in a previous study (Bonnot et al., 2017). Also, *VPT* and *PHO1* orthologous genes were found highly conserved from green algae to land plants. While *NLA* and *PHF* genes, involved in the regulation of PHT transporters, were present only in the streptophyte lineages (*Figure 1a*). The establishment of this regulatory mechanism could be linked to the process of land colonization by plants, as a possible innovation in the regulation of Pi uptake in terrestrial environments. Probably, this innovation occurred to avoid the Pi toxicity in soil, since the concentration of available Pi on land is higher than in aquatic environments.

### 3) Low redundancy in the Marchantia genetic response to low Pi availability

As in other reports, our evolutionary analysis revealed a low genetic redundancy on key gene families which are involved in Pi sensing, lipid turnover and Pi transport. For example, single copy genes for Mp*SPX*, Mp*SIZ*, Mp*XPL*, Mp*DAG*, Mp*MDG*, Mp*DGDG*, Mp*MGDG*, Mp*SQD*, Mp*VTP* and Mp*NLA* were identified. These findings were relevant since in other plant models, such as *A. thaliana*, those genes belong to families with a large number of members, ranging from 3 to 5. Genetic redundancy complicates the functional characterization of the diverse mechanisms involved in low Pi availability responses in vascular plants. Since in *M. polymorpha* several enzymes involved in glycolipid synthesis are encoded by single genes, this represents an ideal opportunity for future work that can shed light on the evolution of minimal regulatory networks oriented to lipid turnover in response to Pi scarcity. Diverse genes encoding SPX proteins were found as single copy genes, such as Mp*VTP*, Mp*NLA* and Mp*SPX*. In the case of SPX, there are four members in *A. thaliana* conforming a genetic network for the negative regulation of AtPHR1 activity in both high and low Pi conditions and via two independent pathways. However, in *M. polymorpha*, the regulation of MpPHR1 might occur only by the action of MpSPX, which led us to speculate that this protein might be modulating MpPHR1 activity in both high and low Pi conditions (*Figure 6*). In order to test such hypotheses, future and detailed experiments are needed.

### 4) Transcriptional response to cope with limited Pi in *M. polymorpha*

The transcriptional landscape of *M. polymorpha* upon Pi starvation, described in this study, reveals conserved induction of genes involved in several aspects of the response of plants to this stress such as Pi sensing, Pi transport, auxin signaling, OA acid exudation, lipid turnover and ROS signaling, among others. Our work opens several working hypotheses, which include one very interesting gene: Mp*NOX1*, is partially up-regulated under low Pi conditions, suggesting that Pi limitation promotes superoxide radical production via Mp*NOX1* and connects ROS signaling to the morphological response (*Figure 6*), probably to modulate rhizoid development in a manner similar to what occurs in *A. thaliana* root hairs (Foreman et al., 2003). Other interesting genes identified were those encoding CLE peptides, the expression of Mp*CLE2* was up-regulated, whereas Mp*CLE1* was down-regulated under low Pi (Figure 4; *Figure 6*). Previous work showed that ectopic expression of Mp*CLE1* reduces thallus growth area of Marchantia plants grown in sufficient Pi conditions, while loss of function *mpcle1* mutant (Hirakawa et al., 2019) alters the thallus proliferative region and affects the organization of dividing cells. On the other hand, exogenous application of MpCLE2 peptide alters the differentiation of stem cell derived daughters, causing the accumulation of the stem cell pool which, after MpCLE2 removal, leads to dichotomous branching (Hirakawa et al 2020).

Our transcriptional analysis showed that Mp*CLE2* levels are induced upon Pi starvation, while those of Mp*CLE1* are repressed. We hypothesized that, under low Pi conditions, the altered expression of both Mp*CLE1* and Mp*CLE2* may be affecting the proper organization and maintenance of the meristem, which could be reflected in the aberrant development of the thalli observed in plants growing for prolonged times under Pi conditions. Future studies including time-lapse experiments of reporter lines with transcriptional fusions of both Mp*CLE* genes, in contrasting Pi conditions, will be very useful to explore this hypothesis.

Our study not only describes the transcriptional response of *M. polymorpha* upon Pi starvation, but also integrates a genome wide search for the P1BS motive enrichment. Our results show a clear enrichment in P1BS motifs in promoters of DEGs as compared to the nDEGs universe (*Figure 5a,c,e*). Moreover, GO analysis of up-regulated genes harboring the P1BS CRE show subcategories typically related to PSR such as *Pi transport*, *lipase activity* and *ribonuclease activity* (*Figure 5b*). However, there are several other functional categories in the up regulated DEGs with P1BS (*Figure 5b*). One possibility is that MpPHR1, or other MYB-CC TFs could participate in modulating the expression of genes that participate in other biological processes. This hypothesis is in agreement with early reports in *A. thaliana*, where AtPHR1 was shown to be involved in the regulation of genes related to zinc, iron and sculpture homeostasis (Briat et al., 2015; Li and Lan, 2015). Moreover, AtPHR1 controls the expression of some genes involved in the plant immune response, indicating that the genetic network controlled by AtPHR1 is highly complex and responds to several environmental cues (Castrillo et al., 2017).

Another interesting finding from these analyses is that the percentage of promoters with P1BS from the up-regulated DEGs *versus* that of down-regulated DEGs is not so contrasting (*Figure 5a, c and e*). This result is intriguing, despite the number of up-regulated DEGs is higher than that of down-regulated DEGs. We speculate on why a gene that has the P1BS motif in its promoter is down-regulated in response to Pi starvation, one possibility is that, if we consider that MpPHR1 and MpPHR-like TFs are all able to bind the P1BS motif, some of the members of the family may act as repressors over a subset of targets. This type of opposite action of diverse members of a TF family has been shown in *M. polymorpha* for MpARFs (Flores-Sandoval et al., 2015; Kato et al., 2020). Future studies should explore such scenario, since the GO analyses of the down regulated DEGs, containing the P1BS, revealed interesting functional categories such as *Cysteine type endopeptidase inhibitor*, that contain genes which orthologs in other species are expressed in response to diverse plant pathogen and some components of plant immune system, such as papain-type *CYSTEINE ENDOPEPTIDASE 1 AtCEP1, PATHOGEN RESPONSE 1 AtPR1* and *PLANTDEFENSIN* 1.2 *AtPDF1.2* (Höwing et al., 2014; Castrillo et al., 2017). This suggests that *M. polymorpha* might also be prioritizing the nutrition mechanisms over the pathogen defence response, as it occurs in other plant species (Castrillo et al., 2017). Future studies should include the generation loss of function mutants of Mp*PHR1* and *MpPHR-like* genes in order to know the direct targets for each of them, as well as to explore their regulatory roles in response not only to low Pi, but also upon other biotic and abiotic stresses.

Collectively, our morphophysiological, transcriptional and promoter analyses for P1BS, revealed a core of partially conserved responses deployed to cope with low Pi availability in *M. polymorpha*, which were analogous to those displayed by *A. thaliana*. We consider that such conserved adaptations, and the *M. polymorpha* specific traits, should be revised and experimentally tested, case by case, in order to characterize in detail the global strategy of this liverwort to deal with Pi scarce environments.

## Materials and methods

### Evolutionary analysis

In this work we searched for classical genes related to phosphate sensing, root system architecture modification, organic acid exudation and phospholipid turnover along viridiplantae phylogeny. We selected as the query, well known, and previously characterized genes, involved in the response to low Pi availability in *Arabidopsis thaliana* (*Figure 1a*). To identify the orthologous genes in *M. polymorpha* and other 23 representative genomes along the Viridiplantae phylogeny, we performed a search by sequence homology with the software blast v.3.1 (Altschul et al., 1997) used as threshold parameter an e-value 1e-30 for land plants and to green algae genomes an e-value 1e-10. Additionally, a reciprocal blast against Arabidopsis protein database TAIR.ver.10 under the same threshold conditions were carried out. The multiple sequence alignment (MSA) was generated with MAFFT software version 7, on the online website (Katoh & Standley, 2013). After, the MSA was trimmed using TrimAL software version 3, maintaining at least 40% for each original protein alignment (Capella-Gutiérrez et al., 2009). We then selected the best model fit to carry out the phylogenetic inference, software ModelTest was used to analyze the sequences according to ACR criterion (Posada & Crandall, 1998). For the phylogenetic reconstruction was obtained with RAxML software launched on the CIPRES environment (https://www.phylo.org/portal2/home.action) (Miller et al., 2010). The bootstrapping value used for each reconstruction was 1000 and the best tree was plotted in iTOL online tool (https://itol.embl.de/) (Letunik & Bork, 2019).

### Plant material and growth conditions

Gemmaelings from *M. polymorpha* (Tak-1) were sown under Murashige and Skoog (MS) media (1/10), pH 5.5, 1% of Sucrose (W/V) and agar 10 g/L. The plants grown under continuous light in Percival chambers with a light intensity, temperature was 22 °C and humidity around to 60%. To determine the Pi concentration for high and low availability, gemmaelings were exposed to a gradient of K2PO4 (0, 10, 25, 50, 100 and 500 μM) for 21 days. Phenotypic effects were registered by pictures in the stereoscope Zeiss at 7, 14 and 21 days after sown (DAS). For further experiments were determined growth conditions used for low Pi (-Pi) 10 μM and high Pi (+Pi) 500 μM of K2PO4 respectively. Also, for RNAseq experiment we use MS (1/10) liquid media, pH 5.5, 1% of Sucrose (W/V), supplemented with K2PO4 to establish the +Pi and −Pi conditions. The implementation of liquid media for RNAseq sample preparation was due to in solid media thallus carried agar traces in the rhizoids. The plants grow in +Pi conditions for 10 days under continuous light, 22 °C and humidity around to 60%. Then we washed the thallus with distilled water and transferred it to +Pi and −Pi media. After 12, 24 and 150 hours the tissue was collected for RNA extraction.

### *In vivo* phosphatase activity assay

The gemmae from *M. polymorpha* (Tak1) were sown under MS media (1/10) supplemented with 1% of Sucrose (w/v), phytagel 3 g/L, pH 5.5 and displayed on polycarbonate cell growth chambers. Also, the Pi (K2PO4) was supplied to establish the +Pi and −Pi conditions, respectively. To detect the phosphatase activity we added after sterilization ELF-97 reagent to final concentration of 50 μM. Plants were grown in Percival chambers with continuous light, temperature 22 °C and humidity around to 60% for 5 days. The images were taken in the confocal microscope Zeiss LM800, we see the chlorophyll auto-florescence at 610 nm and the ELF-97 signal at 530 nm.

### Free phosphate quantification

To perform the quantification of free Pi we used the malachite green assay (Kanno et al., 2016) modified to work with young thallus. First, we ground the fresh samples (50 mg) in liquid nitrogen, added 500 μL of MES buffer (0.17 M, 100 mM DTT, pH 5.8) to each sample and mixed vigorously. After centrifugation at 12 000 rpm for 10 minutes, the supernatant was collected in a new tube. Then 50 μL of each sample were mixed with 115 μL of acid molybdate solution (17.55 g/L of Ammonium molybdate tetrahydrate, 3M sulfuric acid) and 700 μL of deionized water. Ten minutes later, was added 150 μL of green malachite solution (3.5 g/L polyvinyl alcohol, 0.35 g/L malachite green). The absorbance was read at 610 nm after 2 h. A titration curve was performed in the same condition with stock solution 1 mM KH2PO4 to measure 25, 12.5, 6.25, 3.13, 1.56 and 0 μM respectively. Finally, to calculate the Pi concentration in each sample we used a regression equation and normalized by fresh weight.

### RNA extraction, sequencing and Quantitative real-time qPCR analysis

Samples for RNA extraction were collected and ground in liquid nitrogen. For each sample was added 900 μL of TRIzol reagent and were incubated in ice for several minutes and 5 minutes at room temperature (RT). Previous to centrifugation at 13 000 rpm for 10 min at 4 °C, we add 200 μL of Chloroform/Isoamyl alcohol (24:1) and mix briefly for 2 min at RT. The supernatant was collected in a new tube and was incubated for 10 min at RT with 500 μL of Isopropyl alcohol stored previously at - 20 °C, later was precipitated by centrifugation at 13 000 rpm for 10 min at 4 °C. Finally, the precipitated RNA was washed with ethanol 75% (V/V) stored at −20 °C and was resuspended in DEPC water. Before sequencing, the RNA samples were analyzed with Bioanalizer and RIN values lower than 8 were discharged. Library preparation used TruSeq RNA and sequencing with ILLUMINA next seq 1×75. For qPCR experiments total RNA were extracted and analyzed as described above. We used 5 μg of total RNA for each 20 μL reaction to obtain cDNA for each condition and time. cDNA was synthesized using SuperScript II (Invitrogen) according to manufacturer’s protocol. The expression quantification was determined using SYBR Green PCR master mix (Applied Biosystems) in 10 μL reaction. The levels for each tested transcript were determined using three technical replicates per biological replicate and normalized to Actin transcript levels. The results are represented as [2^(-Δ C_T_)], the primers used in these reactions are enlisted *Supplementary figure S36*.

### Differential expressed genes determination and GO enrichment analysis

The raw data were collected to perform quality analysis with the software FastQC. Also, the reads were processed with Trimmomatic software to remove the adapter contamination (Bolger et al., 2014). Then we upload the filtered data to the Cyverse environment (https://de.cyverse.org/de/) and use the DNA subway tool (https://dnasubway.cyverse.org/) to generate a RNAseq analysis to obtain the differential gene expressed genes. We used the pipeline established for the tuxedo suit, the DEG were selected based on FC (> 1.5 and < −1.5) and FDR < 0.001. Venn diagrams were performed using the online tool (http://bioinformatics.psb.ugent.be/webtools/Venn/). On the other hand the enrichment analysis of gene ontology (GO) terms was carried out for the DEG selected each time. The GO analysis was performed in the online tool *PlantRegMap* (http://plantregmap.gao-lab.org/go.php), we selected the *M. polymorpha* genome to perform the analysis of biological processes categories and the P-value lower than 0.001.

### P1BS search in DEGs

To identify the PHR1 DNA binding site (P1BS) in the cis-regulatory region of differentially expressed genes, we first retrieved a 3 kb fragment upstream of the start codon of all *M. polymorpha* genes detected in our transcriptomes using the v3.1 genome release available in Phytozome 13 (https://phytozome-next.jgi.doe.gov/). Then, we searched for the presence of P1BS in these cis-regulatory regions using HOMER (v4.11, 10-24-2019) (Heinz et al., 2010). For this, we used the P1BS consensus sequence (*GNATATNC*) to make a motif file with the seq2profile.pl tool and then we queried the cis-regulatory regions of all the differentially expressed genes with the findMotifs.pl tool. The upstream regions of non-differentially expressed genes were selected as background for this analysis. Once we had the P1BS occurrences and their sequence, we built a sequence logo that represented the P1BS in *M. polymorpha* with ggseqlogo (Wagih, 2017) in R.

### Analysis of P1BS enrichment

To determine if the P1BS was relatively enriched in the promoter set of differentially expressed genes compared with the promoter set of all the genes in our transcriptome, we used AME (McLeay & Bailey, 2010), a tool from the MEME suite (Bailey et al., 2009). We provided AME with the P1BS motif sequence to search and did two analyses in which we wanted to test for P1BS enrichment: the set of 3259 promoters with at least 1 P1BS obtained from the initial P1BS search with HOMER, and the 10783 promoters belonging to non-differentially expressed genes. For both analyses we utilized the 16240 promoters available in the *Marchantia polymorpha* v3.1 genome release (https://phytozome.jgi.doe.gov) as a control sequence set. We selected an average odds score as a sequence scoring method and the Fisher’s exact test as the motif enrichment test.

### Supplementary Materials and RNAseq datasets accession number will be available after acceptance

## References

Alcaraz, L. D., Olmedo, G., Bonilla, G., Cerritos, R., Hernández, G., Cruz, A.,… & López, V. (2008). The genome of Bacillus coahuilensis reveals adaptations essential for survival in the relic of an ancient marine environment. Proceedings of the National Academy of Sciences, 105(15), 5803–5808.

Altschul, S. F., Madden, T. L., Schäffer, A. A., Zhang, J., Zhang, Z., Miller, W., & Lipman, D. J. (1997). Gapped BLAST and PSI-BLAST: a new generation of protein database search programs. Nucleic acids research, 25(17), 3389–3402.

Andersson, M. X., Stridh, M. H., Larsson, K. E., Liljenberg, C., & Sandelius, A. S. (2003). Phosphate-deficient oat replaces a major portion of the plasma membrane phospholipids with the galactolipid digalactosyldiacylglycerol. FEBS letters, 537(1-3), 128–132.

Balzergue, C., Dartevelle, T., Godon, C., Laugier, E., Meisrimler, C., Teulon, J. M.,… & Müller, J. (2017). Low phosphate activates STOP1-ALMT1 to rapidly inhibit root cell elongation. Nature communications, 8, 15300.

Bariola, P. A., Howard, C. J., Taylor, C. B., Verburg, M. T., Jaglan, V. D., & Green, P. J. (1994). The Arabidopsis ribonuclease gene RNS1 is tightly controlled in response to phosphate limitation. The Plant Journal, 6(5), 673–685.

Berland, H., Albert, N. W., Stavland, A., Jordheim, M., McGhie, T. K., Zhou, Y.,… & Davies, K. M. (2019). Auronidins are a previously unreported class of flavonoid pigments that challenges when anthocyanin biosynthesis evolved in plants. Proceedings of the National Academy of Sciences, 116(40), 20232–20239.

Bieleski, R. L. (1973). Phosphate pools, phosphate transport, and phosphate availability. Annual review of plant physiology, 24(1), 225–252.

Bindea, G., Mlecnik, B., Hackl, H., Charoentong, P., Tosolini, M., Kirilovsky, A.,… & Galon, J. (2009). ClueGO: a Cytoscape plug-in to decipher functionally grouped gene ontology and pathway annotation networks. Bioinformatics, 25(8), 1091–1093.

Bolger, A. M., Lohse, M., & Usadel, B. (2014). Trimmomatic: a flexible trimmer for Illumina sequence data. Bioinformatics, 30(15), 2114–2120.

Bonnot, C., Proust, H., Pinson, B., Colbalchini, F. P., Lesly-Veillard, A., Breuninger, H.,… & Dolan, L. (2017). Functional PTB phosphate transporters are present in streptophyte algae and early diverging land plants. New Phytologist, 214(3), 1158–1171.

Bowman, J. L., Kohchi, T., Yamato, K. T., Jenkins, J., Shu, S., Ishizaki, K.,… & Adam, C. (2017). Insights into land plant evolution garnered from the Marchantia polymorpha genome. Cell, 171(2), 287–304.

Briat, J. F., Rouached, H., Tissot, N., Gaymard, F., & Dubos, C. (2015). Integration of P, S, Fe, and Zn nutrition signals in Arabidopsis thaliana: potential involvement of PHOSPHATE STARVATION RESPONSE 1 (PHR1). Frontiers in plant science, 6, 290.

Bustos, R., Castrillo, G., Linhares, F., Puga, M. I., Rubio, V., Pérez-Pérez, J.,… & Paz-Ares, J. (2010). A central regulatory system largely controls transcriptional activation and repression responses to phosphate starvation in Arabidopsis. PLoS genetics, 6(9).

Calderón-Vázquez, C., Sawers, R. J., & Herrera-Estrella, L. (2011). Phosphate deprivation in maize: genetics and genomics. Plant physiology, 156(3), 1067–1077.

Capella-Gutiérrez, S., Silla-Martínez, J. M., & Gabaldón, T. (2009). trimAl: a tool for automated alignment trimming in large-scale phylogenetic analyses. Bioinformatics, 25(15), 1972–1973.

Carol, R. J., Takeda, S., Linstead, P., Durrant, M. C., Kakesova, H., Derbyshire, P.,… & Dolan, L. (2005). A RhoGDP dissociation inhibitor spatially regulates growth in root hair cells. Nature, 438(7070), 1013–1016.

Castrillo, G., Teixeira, P. J. P. L., Paredes, S. H., Law, T. F., de Lorenzo, L., Feltcher, M. E.,… & Paz-Ares, J. (2017). Root microbiota drive direct integration of phosphate stress and immunity. Nature, 543(7646), 513–518.

Creelman, R. A., & Mullet, J. E. (1997). Biosynthesis and action of jasmonates in plants. Annual review of plant biology, 48(1), 355–381.

Cruz-Ramírez, A., Oropeza-Aburto, A., Razo-Hernández, F., Ramírez-Chávez, E., & Herrera-Estrella, L. (2006). Phospholipase DZ2 plays an important role in extraplastidic galactolipid biosynthesis and phosphate recycling in Arabidopsis roots. Proceedings of the National Academy of Sciences, 103(17), 6765–6770.

Daram, P., Brunner, S., Rausch, C., Steiner, C., Amrhein, N., & Bucher, M. (1999). Pht2; 1 encodes a low-affinity phosphate transporter from Arabidopsis. The Plant Cell, 11(11), 2153–2166

Del Pozo, J. C., Allona, I., Rubio, V., Leyva, A., De La Peña, A., Aragoncillo, C., & Paz-Ares, J. (1999). A type 5 acid phosphatase gene from Arabidopsis thaliana is induced by phosphate starvation and by some other types of phosphate mobilising/oxidative stress conditions. The Plant Journal, 19(5), 579–589.

Delwiche, C. F., & Cooper, E. D. (2015). The evolutionary origin of a terrestrial flora. Current Biology, 25(19), R899–R910.

Dipp-Álvarez, M., & Cruz-Ramírez, A. (2019). A phylogenetic study of the ANT Family points to a preANT gene as the ancestor of basal and euANT transcription factors in land plants. Frontiers in plant science, 10, 17.

Duan, K., Yi, K., Dang, L., Huang, H., Wu, W., & Wu, P. (2008). Characterization of a sub-family of Arabidopsis genes with the SPX domain reveals their diverse functions in plant tolerance to phosphorus starvation. The Plant Journal, 54(6), 965–975.

Essigmann, B., Güler, S., Narang, R. A., Linke, D., & Benning, C. (1998). Phosphate availability affects the thylakoid lipid composition and the expression of SQD1, a gene required for sulfolipid biosynthesis in Arabidopsis thaliana. Proceedings of the National Academy of Sciences, 95(4), 1950–1955.

Flores-Sandoval, E., Eklund, D. M., & Bowman, J. L. (2015). A simple auxin transcriptional response system regulates multiple morphogenetic processes in the liverwort Marchantia polymorpha. PLoS genetics, 11(5).

Franco-Zorrilla, J. M., Martín, A. C., Leyva, A., & Paz-Ares, J. (2005). Interaction between phosphate-starvation, sugar, and cytokinin signaling in Arabidopsis and the roles of cytokinin receptors CRE1/AHK4 and AHK3. Plant physiology, 138(2), 847–857.

Franco-Zorrilla, J. M., Martin, A. C., Solano, R., Rubio, V., Leyva, A., & Paz-Ares, J. (2002). Mutations at CRE1 impair cytokinin-induced repression of phosphate starvation responses in Arabidopsis. The Plant Journal, 32(3), 353–360.

Gargouri, M., Park, J. J., Holguin, F. O., Kim, M. J., Wang, H., Deshpande, R. R.,… & Gang, D. R. (2015). Identification of regulatory network hubs that control lipid metabolism in Chlamydomonas reinhardtii. Journal of experimental botany, 66(15), 4551–4566.

González, E., Solano, R., Rubio, V., Leyva, A., & Paz-Ares, J. (2005). PHOSPHATE TRANSPORTER TRAFFIC FACILITATOR1 is a plant-specific SEC12-related protein that enables the endoplasmic reticulum exit of a high-affinity phosphate transporter in Arabidopsis. The Plant Cell, 17(12), 3500–3512.

González-Muñoz, E., Avendaño-Vázquez, A. O., Montes, R. A. C., de Folter, S., Andrés-Hernández, L., Abreu-Goodger, C., & Sawers, R. J. (2015). The maize (Zea mays ssp. mays var. B73) genome encodes 33 members of the purple acid phosphatase family. Frontiers in plant science, 6, 341.

Guo, B., Irigoyen, S., Fowler, T. B., & Versaw, W. K. (2008A). Differential expression and phylogenetic analysis suggest specialization of plastid-localized members of the PHT4 phosphate transporter family for photosynthetic and heterotrophic tissues. Plant signaling & behavior, 3(10), 784–790.

Guo, B., Jin, Y., Wussler, C., Blancaflor, E. B., Motes, C. M., & Versaw, W. K. (2008B). Functional analysis of the Arabidopsis PHT4 family of intracellular phosphate transporters. New Phytologist, 177(4), 889–898.

Gutiérrez-Alanís, D., Ojeda-Rivera, J. O., Yong-Villalobos, L., Cárdenas-Torres, L., & Herrera-Estrella, L. (2018). Adaptation to phosphate scarcity: Tips from arabidopsis roots. Trends in plant science, 23(8), 721–730.

Gutiérrez-Alanís, D., Yong-Villalobos, L., Jiménez-Sandoval, P., Alatorre-Cobos, F., Oropeza-Aburto, A., Mora-Macías, J.,… & Herrera-Estrella, L. (2017). Phosphate starvation-dependent iron mobilization induces CLE14 expression to trigger root meristem differentiation through CLV2/PEPR2 signaling. Developmental cell, 41(5), 555–570.

Han, G. Z. (2017). Evolution of jasmonate biosynthesis and signaling mechanisms. Journal of experimental botany, 68(6), 1323–1331.

Heinz S, Benner C, Spann N, Bertolino E et al. (2010) Simple Combinations of Lineage-Determining Transcription Factors Prime cis-Regulatory Elements Required for Macrophage and B Cell Identities. Mol Cell.

Hidayati, N. A., Yamada-Oshima, Y., Iwai, M., Yamano, T., Kajikawa, M., Sakurai, N.,… & Shimojima, M. (2019). Lipid remodeling regulator 1 (LRL 1) is differently involved in the phosphorus-depletion response from PSR 1 in Chlamydomonas reinhardtii. The Plant Journal, 100(3), 610–626.

Hinsinger, P. (2001). Bioavailability of soil inorganic P in the rhizosphere as affected by root-induced chemical changes: a review. Plant and soil, 237(2), 173–195.

Hirakawa, Y., Uchida, N., Yamaguchi, Y. L., Tabata, R., Ishida, S., Ishizaki, K.,… & Bowman, J. L. (2019). Control of proliferation in the haploid meristem by CLE peptide signaling in Marchantia polymorpha. PLoS genetics, 15(3), e1007997.

Höwing, T., Huesmann, C., Hoefle, C., Nagel, M. K., Isono, E., Huckelhoven, R., & Gietl, C. (2014). Endoplasmic reticulum KDEL-tailed cysteine endopeptidase 1 of Arabidopsis (AtCEP1) is involved in pathogen defense. Frontiers in plant science, 5, 58.

Huang, K. L., Ma, G. J., Zhang, M. L., Xiong, H., Wu, H., Zhao, C. Z.,… & Li, X. B. (2018). The ARF7 and ARF19 transcription factors positively regulate PHOSPHATE STARVATION RESPONSE1 in Arabidopsis roots. Plant physiology, 178(1), 413–427.

Huang, Z., Terpetschnig, E., You, W., & Haugland, R. P. (1992). 2-(2’-Phosphoryloxyphenyl)-4 (3H)-quinazolinone derivatives as fluorogenic precipitating substrates of phosphatases. Analytical biochemistry, 207(1), 32–39.

Ishizaki, K. (2017). Evolution of land plants: insights from molecular studies on basal lineages. Bioscience, biotechnology, and biochemistry, 81(1), 73–80.

Jia, F., Wan, X., Zhu, W., Sun, D., Zheng, C., Liu, P., & Huang, J. (2015). Overexpression of mitochondrial phosphate transporter 3 severely hampers plant development through regulating mitochondrial function in Arabidopsis. PLoS One, 10(6), e0129717.

Jiang, C., Gao, X., Liao, L., Harberd, N. P., & Fu, X. (2007). Phosphate starvation root architecture and anthocyanin accumulation responses are modulated by the gibberellin-DELLA signaling pathway in Arabidopsis. Plant physiology, 145(4), 1460–1470.

Kanno, S., Arrighi, J. F., Chiarenza, S., Bayle, V., Berthomé, R., Péret, B.,… & Thibaud, M. C. (2016). A novel role for the root cap in phosphate uptake and homeostasis. Elife, 5, e14577.

Kato, H., Ishizaki, K., Kouno, M., Shirakawa, M., Bowman, J. L., Nishihama, R., & Kohchi, T. (2015). Auxin-mediated transcriptional system with a minimal set of components is critical for morphogenesis through the life cycle in *Marchantia polymorpha*. PLoS genetics, 11(5).

Kato, H., Mutte, S. K., Suzuki, H., Crespo, I., Das, S., Radoeva, T.,… & Lindhoud, S. (2020). Design principles of a minimal auxin response system. Nature Plants, 6(5), 473–482.

Katoh, K., & Standley, D. M. (2013). MAFFT multiple sequence alignment software version 7: improvements in performance and usability. Molecular biology and evolution, 30(4), 772–780.

Kieber, J. J., & Schaller, G. E. (2018). Cytokinin signaling in plant development. Development, 145(4), dev149344.

Kochian, L. V., Piñeros, M. A., Liu, J., & Magalhaes, J. V. (2015). Plant adaptation to acid soils: the molecular basis for crop aluminum resistance. Annual Review of Plant Biology, 66, 571–598.

Kubo, H., Nozawa, S., Hiwatashi, T., Kondou, Y., Nakabayashi, R., Mori, T.,… & Ishizaki, K. (2018). Biosynthesis of riccionidins and marchantins is regulated by R2R3-MYB transcription factors in Marchantia polymorpha. Journal of plant research, 131(5), 849–864.

Letunic, I., & Bork, P. (2019). Interactive Tree Of Life (iTOL) v4: recent updates and new developments. Nucleic acids research, 47(W1), W256–W259.

Li, D., Zhu, H., Liu, K., Liu, X., Leggewie, G., Udvardi, M., & Wang, D. (2002). Purple acid phosphatases of Arabidopsis thaliana comparative analysis and differential regulation by phosphate deprivation. Journal of Biological Chemistry, 277(31), 27772–27781.

Li, W., & Lan, P. (2015). Genome-wide analysis of overlapping genes regulated by iron deficiency and phosphate starvation reveals new interactions in Arabidopsis roots. BMC research notes, 8(1), 555.

Liu, J., Magalhaes, J. V., Shaff, J., & Kochian, L. V. (2009). Aluminum-activated citrate and malate transporters from the MATE and ALMT families function independently to confer Arabidopsis aluminum tolerance. The Plant Journal, 57(3), 389–399.

Liu, J., Yang, L., Luan, M., Wang, Y., Zhang, C., Zhang, B.,… & Luan, S. (2015). A vacuolar phosphate transporter essential for phosphate homeostasis in Arabidopsis. Proceedings of the National Academy of Sciences, 112(47), E6571–E6578.

López-Bucio, J., Cruz-Ramirez, A., & Herrera-Estrella, L. (2003). The role of nutrient availability in regulating root architecture. Current opinion in plant biology, 6(3), 280–287.

Luan, M., Zhao, F., Han, X., Sun, G., Yang, Y., Liu, J.,… & Luan, S. (2019). Vacuolar phosphate transporters contribute to systemic phosphate homeostasis vital for reproductive development in Arabidopsis. Plant physiology, 179(2), 640–655.

Lv, Q., Zhong, Y., Wang, Y., Wang, Z., Zhang, L., Shi, J.,… & Wu, P. (2014). SPX4 negatively regulates phosphate signaling and homeostasis through its interaction with PHR2 in rice. The Plant Cell, 26(4), 1586–1597.

Lynch, J. (1995). Root architecture and plant productivity. Plant physiology, 109(1), 7.

Ma, Z., Bielenberg, D. G., Brown, K. M., & Lynch, J. P. (2001). Regulation of root hair density by phosphorus availability in Arabidopsis thaliana. Plant, Cell & Environment, 24(4), 459–467.

Mano, Y., & Nemoto, K. (2012). The pathway of auxin biosynthesis in plants. Journal of experimental Botany, 63(8), 2853–2872.

Marschner, H. (2011). Marschner’s mineral nutrition of higher plants. Academic press.

Mehra, P., Pandey, B. K., & Giri, J. (2017). Improvement in phosphate acquisition and utilization by a secretory purple acid phosphatase (OsPAP21b) in rice. Plant Biotechnology Journal, 15(8), 1054–1067.

Miller, M. A., Pfeiffer, W., & Schwartz, T. (2010, November). Creating the CIPRES Science Gateway for inference of large phylogenetic trees. In 2010 gateway computing environments workshop (GCE) (pp. 1–8). Ieee.

Misson, J., Raghothama, K. G., Jain, A., Jouhet, J., Block, M. A., Bligny, R.,… & Doumas, P. (2005). A genome-wide transcriptional analysis using Arabidopsis thaliana Affymetrix gene chips determined plant responses to phosphate deprivation. Proceedings of the National Academy of Sciences, 102(33), 11934–11939.

Mittler, R., Vanderauwera, S., Suzuki, N., Miller, G. A. D., Tognetti, V. B., Vandepoele, K.,… & Van Breusegem, F. (2011). ROS signaling: the new wave?. Trends in plant science, 16(6), 300–309.

Mok, D. W., & Mok, M. C. (2001). Cytokinin metabolism and action. Annual review of plant biology, 52(1), 89–118.

Mora-Macías, J., Ojeda-Rivera, J. O., Gutiérrez-Alanís, D., Yong-Villalobos, L., Oropeza-Aburto, A., Raya-González, J.,… & Herrera-Estrella, L. (2017). Malate-dependent Fe accumulation is a critical checkpoint in the root developmental response to low phosphate. Proceedings of the National Academy of Sciences, 114(17), E3563–E3572.

Morcuende, R., Bari, R., Gibon, Y., Zheng, W., Pant, B. D., BlÄSing, O.,… & Scheible, W. R. (2007). Genome-wide reprogramming of metabolism and regulatory networks of Arabidopsis in response to phosphorus. Plant, Cell & Environment, 30(1), 85–112.

Moseley, J., & Grossman, A. R. (2009). Phosphate metabolism and responses to phosphorus deficiency. In The Chlamydomonas Sourcebook (pp. 189–215). Academic Press.

Müller, J., Toev, T., Heisters, M., Teller, J., Moore, K. L., Hause, G.,… & Abel, S. (2015). Iron-dependent callose deposition adjusts root meristem maintenance to phosphate availability. Developmental cell, 33(2), 216–230.

Nakamura, Y., Koizumi, R., Shui, G., Shimojima, M., Wenk, M. R., Ito, T., & Ohta, H. (2009). Arabidopsis lipins mediate eukaryotic pathway of lipid metabolism and cope critically with phosphate starvation. Proceedings of the national academy of sciences, 106(49), 20978–20983.

Nussaume, L., Kanno, S., Javot, H., Marin, E., Nakanishi, T. M., & Thibaud, M. C. (2011). Phosphate import in plants: focus on the PHT1 transporters. Frontiers in plant science, 2, 83.

Osorio, M. B., Ng, S., Berkowitz, O., De Clercq, I., Mao, C., Shou, H.,… & Jost, R. (2019). SPX4 Acts on PHR1-Dependent and-Independent Regulation of Shoot Phosphorus Status in Arabidopsis. Plant physiology, pp-00594.

Pant, B. D., Burgos, A., Pant, P., Cuadros-Inostroza, A., Willmitzer, L., & Scheible, W. R. (2015). The transcription factor PHR1 regulates lipid remodeling and triacylglycerol accumulation in Arabidopsis thaliana during phosphorus starvation. Journal of experimental botany, 66(7), 1907–1918.

Park, B. S., Seo, J. S., & Chua, N. H. (2014). NITROGEN LIMITATION ADAPTATION recruits PHOSPHATE2 to target the phosphate transporter PT2 for degradation during the regulation of Arabidopsis phosphate homeostasis. The Plant Cell, 26(1), 454–464.

Péret, B., Clément, M., Nussaume, L., & Desnos, T. (2011). Root developmental adaptation to phosphate starvation: better safe than sorry. Trends in plant science, 16(8), 442–450.

Pérez-Torres, C. A., López-Bucio, J., Cruz-Ramírez, A., Ibarra-Laclette, E., Dharmasiri, S., Estelle, M., & Herrera-Estrella, L. (2008). Phosphate availability alters lateral root development in Arabidopsis by modulating auxin sensitivity via a mechanism involving the TIR1 auxin receptor. The Plant Cell, 20(12), 3258–3272.

Pires, N. D., & Dolan, L. (2012). Morphological evolution in land plants: new designs with old genes. Philosophical Transactions of the Royal Society B: Biological Sciences, 367(1588), 508–518.

Poirier, Y., Thoma, S., Somerville, C., & Schiefelbein, J. (1991). Mutant of Arabidopsis deficient in xylem loading of phosphate. Plant physiology, 97(3), 1087–1093.

Posada, D., & Crandall, K. A. (1998). Modeltest: testing the model of DNA substitution. Bioinformatics (Oxford, England), 14(9), 817–818.

Proust, H., Honkanen, S., Jones, V. A., Morieri, G., Prescott, H., Kelly, S.,… & Dolan, L. (2016). RSL class I genes controlled the development of epidermal structures in the common ancestor of land plants. Current Biology, 26(1), 93–99.

Puga, M. I., Mateos, I., Charukesi, R., Wang, Z., Franco-Zorrilla, J. M., de Lorenzo, L.,… & Leyva, A. (2014). SPX1 is a phosphate-dependent inhibitor of Phosphate Starvation Response 1 in Arabidopsis. Proceedings of the National Academy of Sciences, 111(41), 14947–14952.

Puga, M. I., Rojas-Triana, M., de Lorenzo, L., Leyva, A., Rubio, V., & Paz-Ares, J. (2017). Novel signals in the regulation of Pi starvation responses in plants: facts and promises. Current opinion in plant biology, 39, 40–49.

Quisel, J. D., Wykoff, D. D., & Grossman, A. R. (1996). Biochemical characterization of the extracellular phosphatases produced by phosphorus-deprived Chlamydomonas reinhardtii. Plant physiology, 111(3), 839–848.

Raghothama, K. G. (1999). Phosphate acquisition. Annual review of plant biology, 50(1), 665–693.

Raya-González, J., Pelagio-Flores, R., & López-Bucio, J. (2012). The jasmonate receptor COI1 plays a role in jasmonate-induced lateral root formation and lateral root positioning in Arabidopsis thaliana. Journal of plant physiology, 169(14), 1348–1358.

Reinhard, C. T., Planavsky, N. J., Gill, B. C., Ozaki, K., Robbins, L. J., Lyons, T. W.,… & Konhauser, K. O. (2017). Evolution of the global phosphorus cycle. Nature, 541(7637), 386–389.

Robert McLeay and Timothy L. Bailey (2010) Motif Enrichment Analysis: A unified framework and method evaluation. BMC Bioinformatics, 11:165 doi:10.1186/1471-2105-11-165.

Rubio, V., Linhares, F., Solano, R., Martín, A. C., Iglesias, J., Leyva, A., & Paz-Ares, J. (2001). A conserved MYB transcription factor involved in phosphate starvation signaling both in vascular plants and in unicellular algae. Genes & development, 15(16), 2122–2133.

Sakakibara, H. (2006). Cytokinins: activity, biosynthesis, and translocation. Annu. Rev. Plant Biol., 57, 431–449.

Sanderson, M. J. (2003). Molecular data from 27 proteins do not support a Precambrian origin of land plants. American Journal of Botany, 90(6), 954–956.

Secco, D., Jabnoune, M., Walker, H., Shou, H., Wu, P., Poirier, Y., & Whelan, J. (2013). Spatio-temporal transcript profiling of rice roots and shoots in response to phosphate starvation and recovery. The Plant Cell, 25(11), 4285–4304.

Shimamura, M. (2016). Marchantia polymorpha: taxonomy, phylogeny and morphology of a model system. Plant and Cell Physiology, 57(2), 230–256.

Shimogawara, K., Wykoff, D. D., Usuda, H., & Grossman, A. R. (1999). Chlamydomonas reinhardtii mutants abnormal in their responses to phosphorus deprivation. Plant physiology, 120(3), 685–694.

Shull, C. (1925). Phosphorus and the early maturity of plants. Trans. Illinois State Acad. Sci, 18, 165–71.

Song, L., Yu, H., Dong, J., Che, X., Jiao, Y., & Liu, D. (2016). The molecular mechanism of ethylene-mediated root hair development induced by phosphate starvation. PLoS genetics, 12(7).

Song, S. K., Hofhuis, H., Lee, M. M., & Clark, S. E. (2008). Key divisions in the early Arabidopsis embryo require POL and PLL1 phosphatases to establish the root stem cell organizer and vascular axis. Developmental cell, 15(1), 98–109.

Svistoonoff, S., Creff, A., Reymond, M., Sigoillot-Claude, C., Ricaud, L., Blanchet, A.,… & Desnos, T. (2007). Root tip contact with low-phosphate media reprograms plant root architecture. Nature genetics, 39(6), 792.

Takabatake, R., Hata, S., Taniguchi, M., Kouchi, H., Sugiyama, T., & Izui, K. (1999). Isolation and characterization of cDNAs encoding mitochondrial phosphate transporters in soybean, maize, rice, and Arabidopsis. Plant molecular biology, 40(3), 479–486.

Thibaud, M. C., Arrighi, J. F., Bayle, V., Chiarenza, S., Creff, A., Bustos, R.,… & Nussaume, L. (2010). Dissection of local and systemic transcriptional responses to phosphate starvation in Arabidopsis. The Plant Journal, 64(5), 775–789.

Timothy L. Bailey, Mikael Bodén, Fabian A. Buske, Martin Frith, Charles E. Grant, Luca Clementi, Jingyuan Ren, Wilfred W. Li, William S. Noble (2009) MEME SUITE: tools for motif discovery and searching. Nucleic Acids Research.

Versaw, W. K., & Harrison, M. J. (2002). A chloroplast phosphate transporter, PHT2; 1, influences allocation of phosphate within the plant and phosphate-starvation responses. The Plant Cell, 14(8), 1751–1766.

Voth, P. D. (1941). Gemmae-cup production in Marchantia polymorpha and its response to calcium deficiency and supply of other nutrients. Botanical Gazette, 103(2), 310–325.

Voth, P D., & Hamner, K. C. (1940). Responses of Marchantia polymorpha to nutrient supply and photoperiod. Botanical Gazette, 102(1), 169–205.

Wagih, Omar (2017) ggseqlogo: a versatile R package for drawing sequence logos. Bioinformatics.

Wang, K. L. C., Li, H., & Ecker, J. R. (2002). Ethylene biosynthesis and signaling networks. The plant cell, 14(suppl 1), S131–S151.

Wang, Z., Ruan, W., Shi, J., Zhang, L., Xiang, D., Yang, C.,… & Shou, H. (2014). Rice SPX1 and SPX2 inhibit phosphate starvation responses through interacting with PHR2 in a phosphate-dependent manner. Proceedings of the National Academy of Sciences, 111(41), 14953–14958.

Wellman, C. H., & Strother, P K. (2015). The terrestrial biota prior to the origin of land plants (embryophytes): a review of the evidence. Palaeontology, 58(4), 601–627.

Wickham H (2016). ggplot2: Elegant Graphics for Data Analysis. Springer-Verlag New York. ISBN 978-3-319-24277-4, https://ggplot2.tidyverse.org.

Wild, R., Gerasimaite, R., Jung, J. Y., Truffault, V., Pavlovic, I., Schmidt, A.,… & Mayer, A. (2016). Control of eukaryotic phosphate homeostasis by inositol polyphosphate sensor domains. Science, 352(6288), 986–990.

Wykoff, D. D., Grossman, A. R., Weeks, D. P., Usuda, H., & Shimogawara, K. (1999). Psr1, a nuclear localized protein that regulates phosphorus metabolism in Chlamydomonas. Proceedings of the National Academy of Sciences, 96(26), 15336–15341.

Xu, L., Zhao, H., Wan, R., Liu, Y., Xu, Z., Tian, W.,… & Dolan, L. (2019). Identification of vacuolar phosphate efflux transporters in land plants. Nature plants, 5(1), 84.

Zeng, H., Wang, G., Zhang, Y., Hu, X., Pi, E., Zhu, Y.,… & Du, L. (2016). Genome-wide identification of phosphate-deficiency-responsive genes in soybean roots by high-throughput sequencing. Plant and soil, 398(1-2), 207–227.

Zhang, Y., Zhang, J., Guo, J., Zhou, F., Singh, S., Xu, X.,… & Huang, C. F. (2019). F-box protein RAE1 regulates the stability of the aluminum-resistance transcription factor STOP1 in Arabidopsis. Proceedings of the National Academy of Sciences, 116(1), 319–327.

Zhao, Y. (2010). Auxin biosynthesis and its role in plant development. Annual review of plant biology, 61, 49–64.

